# Design of Peptide Vaccine for COVID19: CD8+ and CD4+ T cell epitopes from SARS-CoV-2 open-reading-frame protein variants

**DOI:** 10.1101/2021.09.21.461301

**Authors:** Simone Parn, Gabriel Jabbour, Vincent Nguyenkhoa, Sivanesan Dakshanamurthy

**Author notes:** Corresponding Author Dr. Sivanesan Dakshanamurthy, Ph.D, MBA.

## Abstract

Severe acute respiratory syndrome coronavirus 2 (SARS-CoV-2), the causative agent of COVID-19, has challenged public health at an unprecedented scale which has led to a dramatic loss of human life worldwide. To design a protective vaccine against SARS-CoV-2, it is necessary to understand which SARS-CoV-2 specific epitopes can elicit a T cell response and provide protection across a broad population. In this study, PLpro and RdRp, two immunogenic non-structural proteins from an immunodominant gene region ORF1ab, as well as ORF3a and ORF9b are identified as potential vaccine targets against SARS-CoV-2. To select top epitopes for vaccine design, we used various clinical properties, such as antigenicity, allergenicity, toxicity and IFN-y secretion. The analysis of CD8 and CD4 T cell epitopes revealed multiple potential vaccine constructs that cover a high percentage of the world population. We identified 8 immunogenic, antigenic, non-allergenic, non-toxic, stable and IFN-y inducing CD8 proteins for nsp3, 4 for nsp12, 11 for ORF3a and 3 for ORF9b that are common across four lineages of variants of concern – B.1.1.7, P.1, B.1.351 and B.1.617.2, which protect 98.12%, 87.08%, 96.07% and 63.8% of the world population, respectively. We also identified variant specific T cell epitopes that could be useful in targeting each variant strain separately. Including the prediction of mouse MHC affinity towards our top CD8 epitopes, our study revealed a total of 3 immunogenic, antigenic, non-allergenic, non-toxic, stable and IFN-y inducing CD8 epitopes overlapping with 6 antigenic, non-allergenic, non-toxic, stable and IFN-y inducing CD4 epitopes across all four variants of concern which can effectively be utilized in pre-clinical studies. The landscape of SARS-CoV-2 T cell epitopes that we identified can help lead SARS-CoV-2 vaccine development as well as epitope-based peptide vaccine research in the future.

## Introduction

Severe acute respiratory syndrome coronavirus 2 (SARS-CoV-2) is a novel causative agent of coronavirus disease 2019 (COVID-19) that belongs to the family of *Coronaviridae* and the genus of *Betacoronavirus* [1]. First emerged in Wuhan, China, the virus rapidly spread across the world, challenging all aspects of human life. On March 11^th^, 2020, the World Health Organization declared the global emergency a pandemic [2]. To date, there has been more than 222 million COVID-19 cases and over 4.5 million deaths worldwide, which indicate the need for a preventative and effective vaccine [3]. In the recent decades, epitope-based vaccine design has gained more insight due to its time- and cost-effective approach [4]. The vaccine approach is centered around immunoinformatics methods and bioinformatics tools that aid in designing CD4+ and CD8+ T cell epitopes against viral antigens. T cells play an important role in the elimination of virus-infected cells, with CD8+ T cells being critical for the clearance of infected cells, and CD4+ T cells for the promotion of specific B cell antibody production and support of CD8+ T cell response. These immune cells have characteristic T cell receptors (TCR) that interact with major histocompatibility complex I and II molecules located on the surface of antigen-presenting cells (APCs), such as dendritic cells, in order to trigger an immune response against a pathogen [5].

The SARS-CoV-2 genome encodes 4 major structural proteins, spike (S), envelope (E), membrane (M), and nucleocapsid (N), and 6 major non-structural proteins, often referred to as open reading frames (ORFs) that carry vital functions for the transmissibility and survival of the virus. ORF1ab, the largest gene region of the SARS-CoV-2 genome, is highly conserved within coronaviruses and it encodes 16 non-structural proteins (nsp1-16) that are cleaved by two polyproteins, pp1a and pp1ab [6]. In this study, PLpro (nsp3) and RdRP (nsp12), were targeted due to their potential ability to provoke the highest immune response within ORF1ab. Nsp3, the largest multi-domain protein produced by coronaviruses, plays a central role in viral replication, as well as nsp12, which functions as RNA polymerase. ORF3a, the largest accessory protein of 275 amino acids in SARS-CoV-2, also has a significant role in virus pathogenesis and T cell reactivity [7, 8, 9].

In addition to ORF1ab and ORF3a proteins, ORF9b, an alternative open reading frame incorporated into the N gene, will be used as another antiviral target in this study. The 97 amino acid long protein is abundant in SARS-CoV-2 and highly conserved in SARS-like coronaviruses, making it a potential vaccine candidate. Besides this, ORF9b plays a critical role in virus-host interactions [10]. Due to its accumulation early in infection and its interaction with TOM70, a mitochondrial protein essential for RNA sensing adaptor MAVS activation, Orf9b suppresses the innate immune response by blocking type I interferon (IFN-I) [11]. Therefore, we aim to target the interaction site between ORF9b and TOM70 by identifying immunogenic CD4+ and CD8+ T cell epitopes that would aid in vaccine construction.

The emergence of variants of SARS-CoV-2 makes it more challenging to come up with protective vaccine candidates. The spectra of SARS-CoV-2 mutations harbors a wide range of silent and point mutations that impact the structure and function of the proteins, making the virus more transmissible. With the virus constantly changing, some variants of concern (VOC) have been detected, including the B.1.1.7, P.1, B.1.351 and B.1.617.2 lineages that first emerged in the United Kingdom, Brazil, South Africa and India, respectively [12]. Therefore, in this analysis the lineage-defining sequences for each VOC will be utilized to shed light on the most important and prevalent T cell epitopes that would help design the most effective vaccine.

In this study, we aim to identify immunogenic CD4+ and CD8+ T cell epitopes on nsp3, nsp12, ORF3a and ORF9b proteins of SARS-CoV-2 for effective vaccine development. It is hypothesized that CD4+ and CD8+ T cell recognition is largely based on HLA specificity, and binding to multiple HLA allelic variants is important for the amplification of potential immunogenicity. We utilized Immune Epitope Database (IEDB) server to compare several available tools for their accuracy predicting immunogenic T cell epitopes [13]. Predicted T cell epitopes were compared by their match rate to experimentally validated immunogenic SARS-CoV-2 T cell epitopes. This allowed the identification of the best performing tool to predict both MHC-I and MHC-II restricted T-cell epitopes from SARS-CoV-2 proteins with high binding affinity to MHC class I and II alleles. Beyond MHC affinity, epitopes were further refined by predicting immunogenicity, population coverage, antigenicity, allergenicity, toxicity, IFN-y secretion, half-life, GRAVY (Grand average of hydropathicity) and other amino acid physiochemical properties, to fully maximize the quality and efficiency of the designed vaccine. The 3D structures of the final CD8 T cell epitopes were thereafter predicted and docked with their respective HLA alleles to visualize the peptide-MHC interaction.

After predicting immunogenic CD4 and CD8 T cells, we attempted to identify top immunogenic CD8 epitopes (8-11 aa long) within immunogenic CD4 epitopes (15 aa long) that together could be used to provoke a strong and long-lasting immune response against the four variants of concern of SARS-CoV-2. In order to safely test the predicted vaccine constructs in pre-clinical studies, we also predicted mouse MHC affinity to our top CD8 epitopes. Mouse models are useful for the evaluation of antiviral targets as they share features with humans and two mouse strains - BALB/CJ and C57BL/6 have been reported as good models for SARS-CoV-2 [14, 15].

## Materials and Methods

### Retrieval of SARS-CoV-2 sequence

The reference protein sequences of SARS-CoV-2 Wuhan isolate were retrieved from NCBI database using accession number NC_045512. The corresponding protein accession numbers are: NCBI: YP_009725299.1 (NSP3), NCBI: YP_009725307.1 (NSP12), NCBI: YP_009724391.1 (ORF3a) and UniProtKB/Swiss-Prot: P0DTD2.1 (ORF9b).

### Retrieval of non-structural protein 3 (nsp3) MHC class I experimental epitopes

To assess the validity of the predictions in the datasets, predicted epitopes were compared to experimentally identified SARS-CoV-2 T cell epitopes. Since nsp3 is the largest multi-domain protein in SARS-Cov-2, the highest number of experimental epitopes was identified and used in the validation. A total of 169 high quality experimental CD8+ T cell epitopes were retrieved from SARS-CoV-2 T cell epitope database that were previously reported in multiple studies [7, 8, 16-23] (**Supplementary Table 1**) and tested for T cell reactivity. Additional 3 epitopes – ILLLDQALV, WLMWLIINL and CLEASFNYL were identified and selected as highly homological to SARS-CoV-1 genome, having identity of 88.89%, 66.67% and 55.56%, respectively [24].

### CD8+ T cell epitope tool comparisons

The IEDB.org website was used to predict SARS-CoV-2 CD8+ T cell epitopes of the reference nsp3 protein using 6 different tools: NetMHCpan EL 4.1 (IEDB recommended), NetMHCpan BA 4.1, NetMHCpan EL 4.0, NetMHCpan BA 4.0 [25-29], ANN 4.0 [30-36], and IEDB consensus [37]. Only MHC I alleles with a frequency >1% were selected for further processing. The duplicate-free peptides were ordered according to their available attributes, such as percentile rank and IC50 value, and then matched to the experimental epitopes using three levels of stringency: lenient, intermediate and stringent. With the lenient method, a predicted epitope matches an experimental epitope if the predicted epitope is included within the experimental epitope or vice versa. With the intermediate method, two epitopes match if they only differ from in up to one single-character edit (insertion, deletion or substitution). With the stringent method, a predicted epitope is only considered a match if it shares 100% sequence similarity with the experimental epitope. To select the best tool, ROC curves with respective AUCs were generated to compare each tool-attribute combination.

### Retrieval of variant sequences

The lineages of variants of concern, UK B.1.1.7 [38], South African B.1.351 [39], Brazilian P.1 [40] and Indian B.1.617.2 [41], were used to identify the most prevalent mutations for nsp3, nsp12, ORF3a and ORF9b that would be present in at least 75% of the sequences [42] (**Table 1**).

**Table 1.** Most prevalent mutations in four variants of concern (VOC): B.1.351, B.1.617.2, B.1.1.7 and P.1. The point mutations are present in at least 75% of identified sequences.

| **Variant** | **ORF1a (nsp3)** | **ORF1ab (nsp12)** | **ORF3a** |
| --- | --- | --- | --- |
| **South Africa**<br>**(B.1.351)** | K837N | P323L | Q57H, S171L |
| **India (B.1.617.2)** | A488S, P1228L,<br>P1469S | P323L, G671S | S26L |
| **UK (B.1.1.7)** | T183I, A890D,<br>I1412T | P323L | N/A |
| **Brazil (P.1)** | S370L, K977Q | P323L | S253P |

### CD8+ T cell epitope prediction and immunogenicity modeling

Using the best tool-attribute combination, CD8+ T cell epitopes were predicted for nsp3, nsp12, ORF3a and ORF9b variants. Then, IEDB immunogenicity tool was used to predict immunogenicity scores for the top 300 duplicate-free epitopes from each variant [43]. Epitopes with an immunogenicity score >0 were selected for further analysis.

### CD4+ T cell epitope prediction

CD4+ T cell variant epitopes were predicted for each respective protein using IEDB recommended 2.22 [44, 45]. Top 300 duplicate-free epitopes based on rank were selected for further analysis.

### IFN-y epitope prediction

IFN-y plays a center role in viral infections for both innate and adaptive immunity. Therefore, it is important to identify IFN-y inducing epitopes. IFNEpitope server was used to predict IFN-y epitopes [46]. Only epitopes with an IFN-y score >0 were selected.

### Antigenicity, allergenicity and toxicity evaluation

Antigenicity was evaluated by VaxiJen 2.0 [47] server with a threshold of 0.4, allergenicity by AllerTop v2.0 [48] and toxicity by ToxinPred [49]. Peptides that were “probable non-allergenic” and non-toxic were selected.

### Amino acid physiochemical properties

Evaluation of various physicochemical properties is essential for determining the safety and efficacy of the candidate vaccine. Hence, various chemical and physical parameters associated with the vaccine were predicted in this study using ProtParam tool on ExPASy [50]. Instability index, aliphatic index, GRAVY and half-life of top CD8 and CD4 epitopes were assessed. To make sure the predicted peptides are stable and not being degraded, instability index <40 was selected as threshold. The aliphatic index points to thermostability, higher the aliphatic index of a protein, greater is its thermostability. The Grand average hydropathicity (GRAVY) indicates the solubility of the peptide. Finally, the half-life in mammalian reticulocytes in vitro was predicted and half-life >1 hour was selected as a threshold value [51].

### Worldwide human population coverage analysis

The population coverage of HLA class I and II alleles, calculated using the IEDB population coverage tool, computes the percentage of the world population predicted to present our epitopes on their MHC molecules. The analysis covered East Asia, Northeast Asia, South Asia, Southeast Asia, Southwest Asia, Europe, East Africa, West Africa, Central Africa, North Africa, South Africa, West Indies, North America, Central America, South America and Oceania [52].

### Murine model MHC restriction

NetH2pan Server [53] was used to predict CD8 epitopes’ affinity to murine MHC class I (H2) molecules. Two common mouse strains to study COVID-19 vaccines, C57BL/6 and BALB/CJ, were used in mouse HLA affinity predictions. Peptides that were either weak binders (WB) or strong binders (SB) were selected.

### 3D dimensional protein structure prediction

The tertiary structure of the top epitopes was predicted using PEPstrMOD [54]. The 3D structure of the respective HLA alleles which nsp3, nsp12, ORF3a and ORF9b epitopes bind to were downloaded from Protein Database Bank on RCSB.org. The native peptides from the RCSB structures were replaced with our predicted epitopes using PyMol. The epitopes were then docked into their respective MHC molecules using the FlexPepDock server [55].

### Visualization of the top CD8 epitopes in 3D structure

The full-length reference protein sequences of ORF3a, nsp3, nsp12 and ORF9b proteins were retrieved from RCSB.org. The proteins and corresponding top CD8 epitopes were visualized using PyMol (**Supplementary Figures 1a-d**).

## Results

### Workflow of T cell predictions

A schematic presentation of the immunoinformatic methods was proposed for epitope predictions (**Figure 3**). The workflow of this study is illustrated in two main steps: IEDB tool comparison and MHC class I and II epitope predictions. Briefly, the sequences of nsp3, nsp12, ORF3a and ORF9b were retrieved from SARS-CoV-2 to predict CD4 and CD8 epitopes. IEDB tool comparison was carried out to identify the best-performing tool-attribute combination for CD8 epitope predictions. Specifically, the nsp3 protein was selected for analysis due to its highest number of high-quality experimental epitopes. After identification of the best prediction tool, CD8 epitopes were screened through IEDB immunogenicity prediction tool, as well as antigenicity, allergenicity, toxicity, IFN-y secretion and several physiochemical properties predictors to select the most effective vaccine constructs. Similar procedure was carried out for CD4 epitopes, however instead of carrying out tool comparisons, the best tool was selected from literature [56].

**Figure 1.**
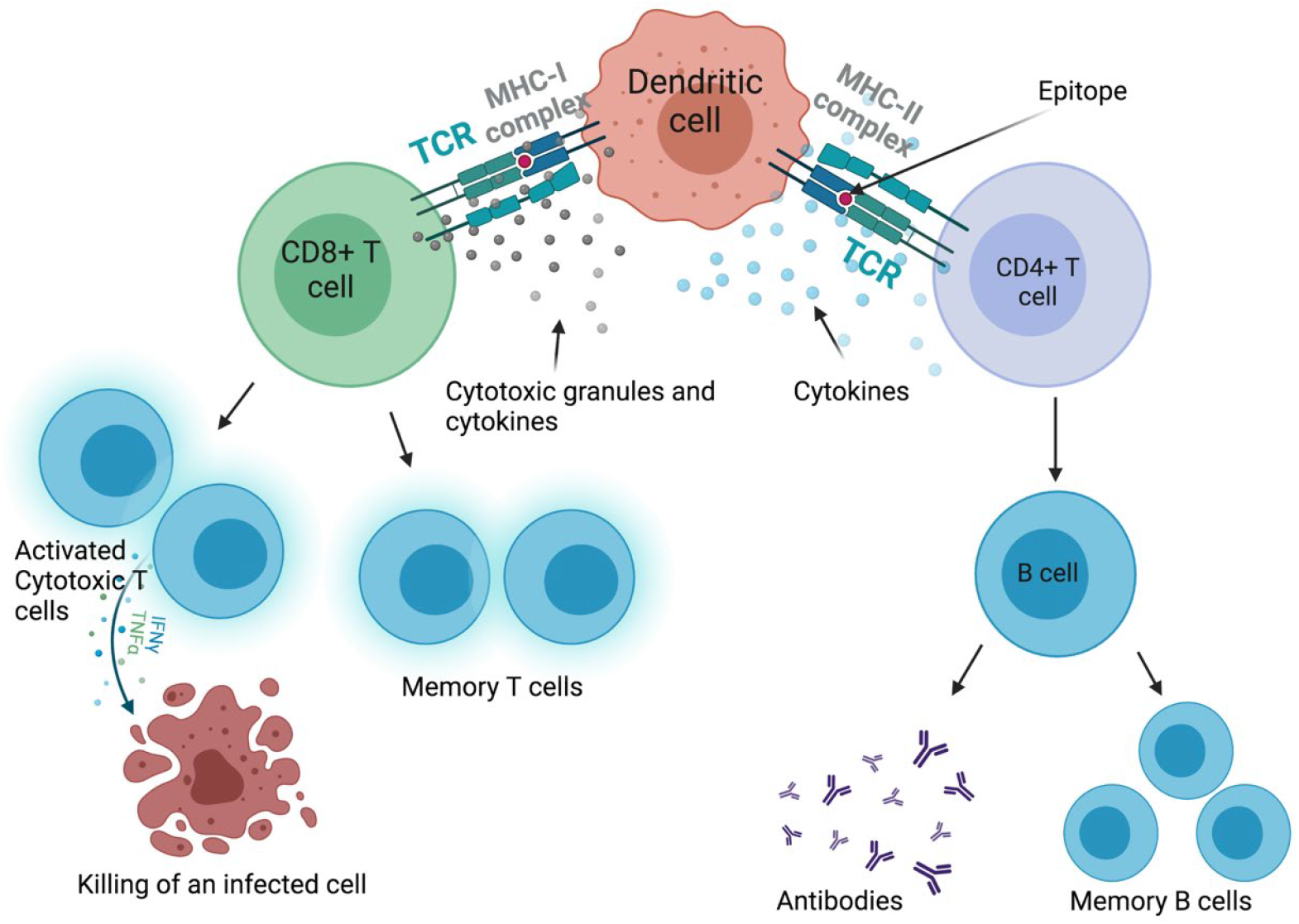
CD4+ and CD8+ T cell activation and response. Interaction between MHC molecules and T cell receptors (TCR) on T cells trigger the activation of CD4+ and CD8+ T cells that lead to the production of memory T and B cells. Cytokines and cytotoxic granules are released in response to a stimulus. The figure was created with BioRender.com.

**Figure 2.**
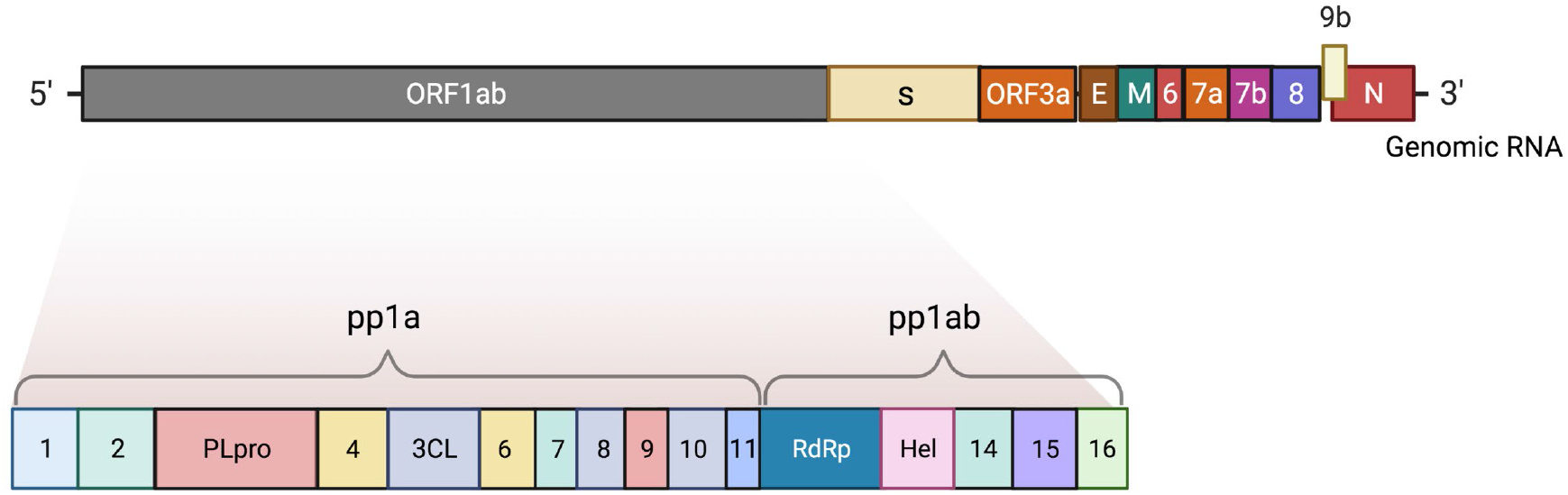
The complete genome of SARS-CoV-2. The 5’ end consists of a large gene region ORF1ab and its non-structural proteins (nsp1-16). The 3’ end compromises the structural proteins (nucleocapsid, membrane, envelope, spike) and other open-reading-frame (ORF) proteins. The figure was created with BioRender.com.

**Figure 3.**
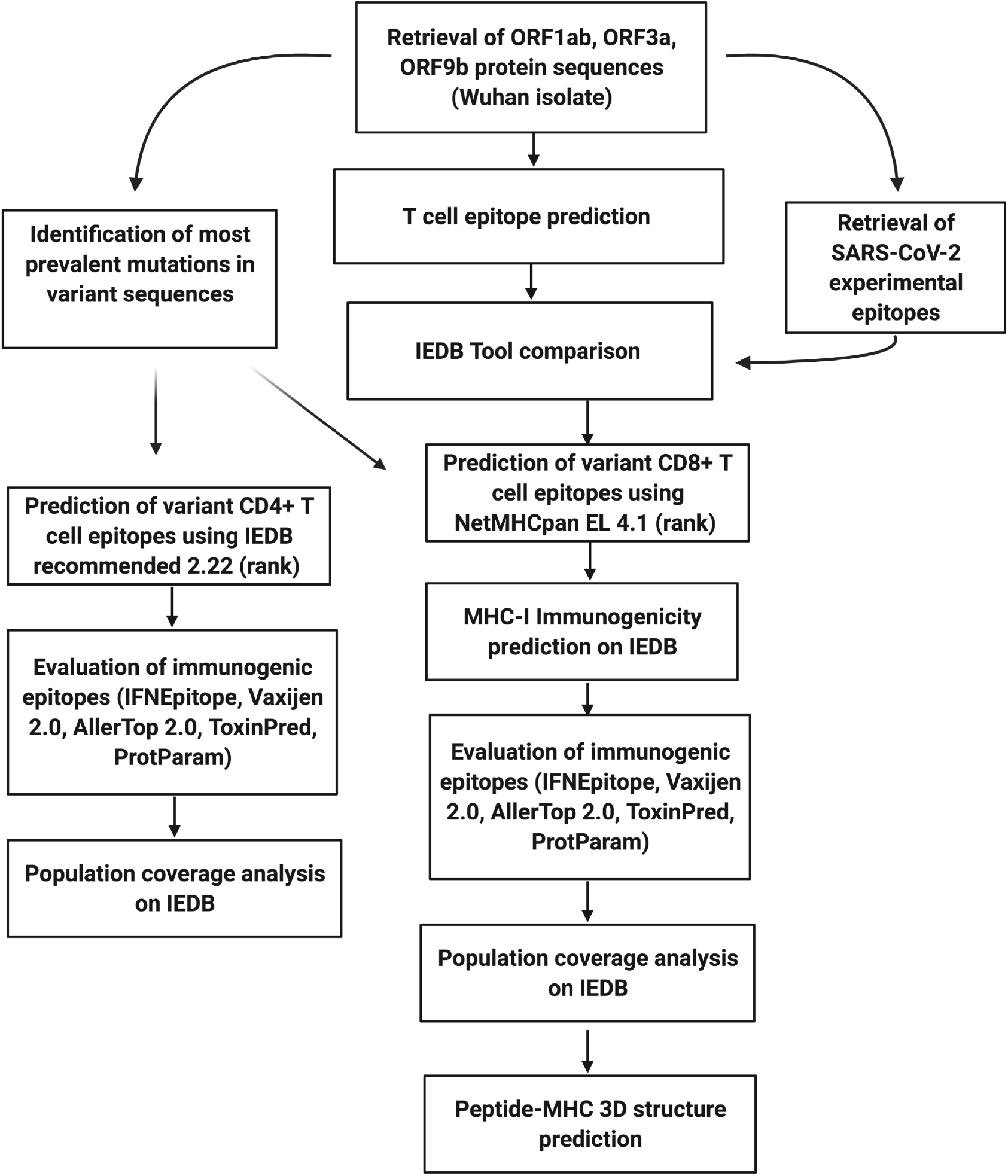
Schematic representation of immunoinformatics identification of T-cell epitopes in SARS-CoV-2. First, protein sequences were retrieved from SARS-CoV-2, which was followed by T cell epitope prediction and IEDB tool comparisons. Variant CD8 and CD4 epitopes were then predicted and analyzed based on their clinical properties. Finally, top common epitopes were used for population coverage and 3D prediction analysis. The figure was created with BioRender.com.

### Tool comparisons for CD8+ T cell binding predictions

The performance of each attribute-tool combination was evaluated by generating receiver operating characteristic curves (ROC) and box plot graphs for each stringency level method (**Figure 4**). A ROC is a probability curve that utilizes Area Under the Curve (AUC) to visualize the performance of a tool-attribute combination [57]. Generally, the higher the AUC value, the better the model is performing. The area under the curve (AUC) values were compared for each stringency level and it was found that IEDB recommended (NetMHCpan EL 4.1 rank) had the highest combined AUC value, meaning its performance was most specific and sensitive (**Figure 4**). Therefore, MHC-I restricted T cell epitopes and their binding affinity were predicted using this tool.

**Figure 4.**
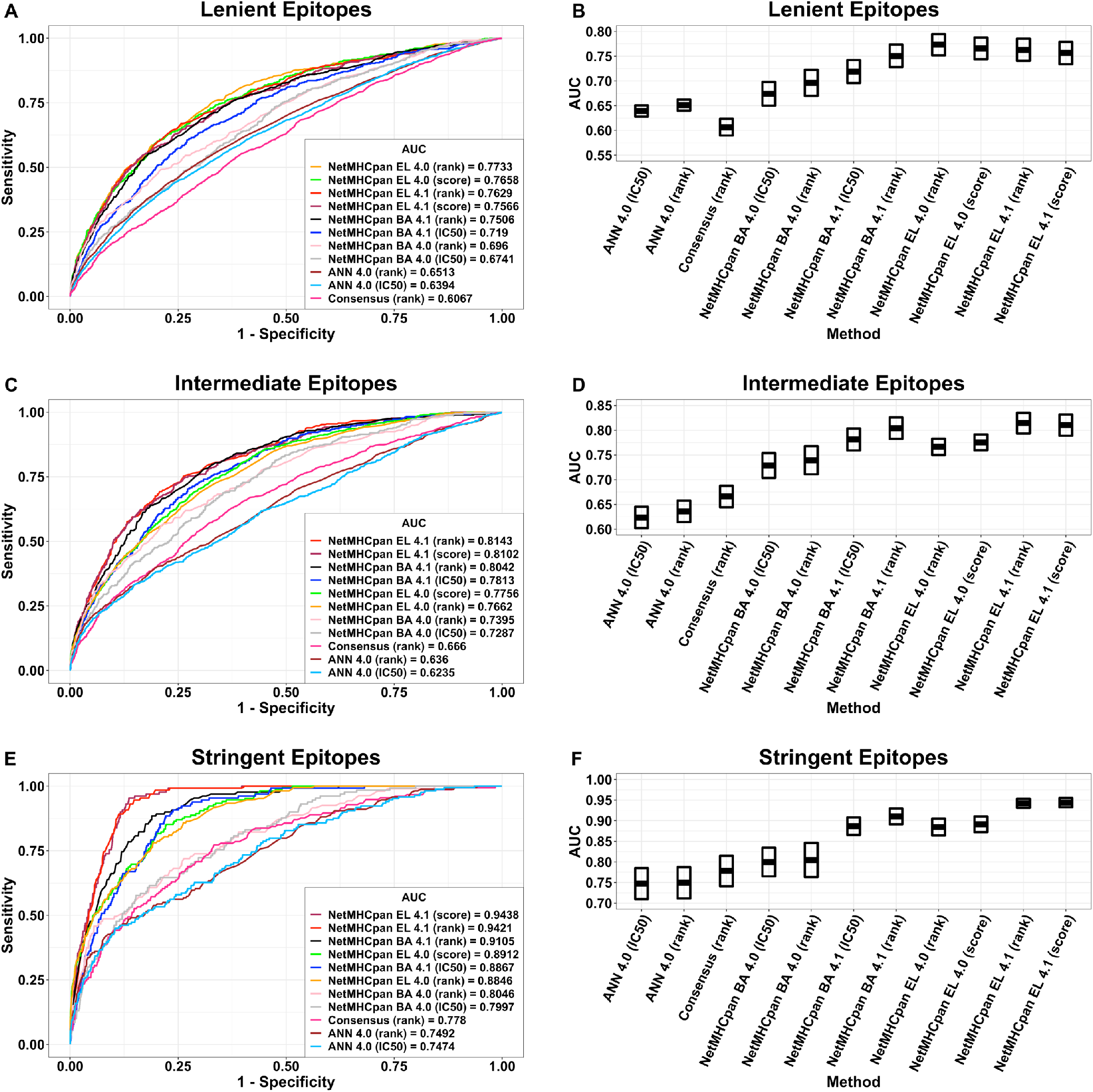
IEDB CD8+ T cell epitope tool comparisons using three levels of stringency: lenient, intermediate and stringent. ROC curves and box plot graphs illustrate the tool-attribute combinations that are useful in the selection of the best tool for CD8 epitope predictions.

### Variant sequences

In order to design the most effective vaccine, it is important to identify the most prevalent mutations across the world. Since the start of the pandemic, four variants of concern (B.1.351, B.1.617.2, B.1.1.7, P.1) have been reported that carry mutations important to virus transmissibility [12]. After filtering through mutations that are presented by at least 75% of the sequences, we were able to identify several nonsynonymous mutations in the nsp3 protein (**Table 1**). On the other hand, the nsp12 protein that is involved in virus replication, was found to have a dominant mutation across all variants of concern (**Table 1**). The point mutation at P323L, first detected in the UK, has been associated with severity of COVID-19 [58]. Also, the amino acid mutation in Q57H of ORF3a has been correlated with severity of the disease [9]. In ORF9b protein, no prevalent mutations were found across the variants of concern.

### CD8+ T cell epitope prediction

Using NetMHCpan EL 4.1 based on rank, MHC-I immunogenicity predictor, IFNEpitope, VaxiJen 2.0, AllerTop 2.0, ToxinPred and ProtParam, only immunogenic, antigenic, IFN-y inducing, non-allergenic, non-toxic and stable CD8 epitopes were selected. We obtained 8 nsp3, 4 nsp12, 11 ORF3a and 3 ORF9b CD8 epitopes, common across all four variants, 1 nsp3 epitope specific to the UK variant and 4 ORF3a epitopes specific to the Indian variant (**Table 2**). While IFNEpitope server is mostly designed for MHC class II restricted epitopes, the MHC class I predictions were also made without the IFN-y filter. We were able to obtain 21 common nsp3, 19 common nsp12, 36 common ORF3a and 5 common ORF9b CD8 epitopes across VOC, 2 nsp3 epitopes specific to the UK variant, 2 nsp12 epitopes specific to the Indian, the UK, Brazilian and South African variants, and 7 ORF3a epitopes specific to the Indian variant (**Supplementary Tables 2-6**).

**Table 2.**
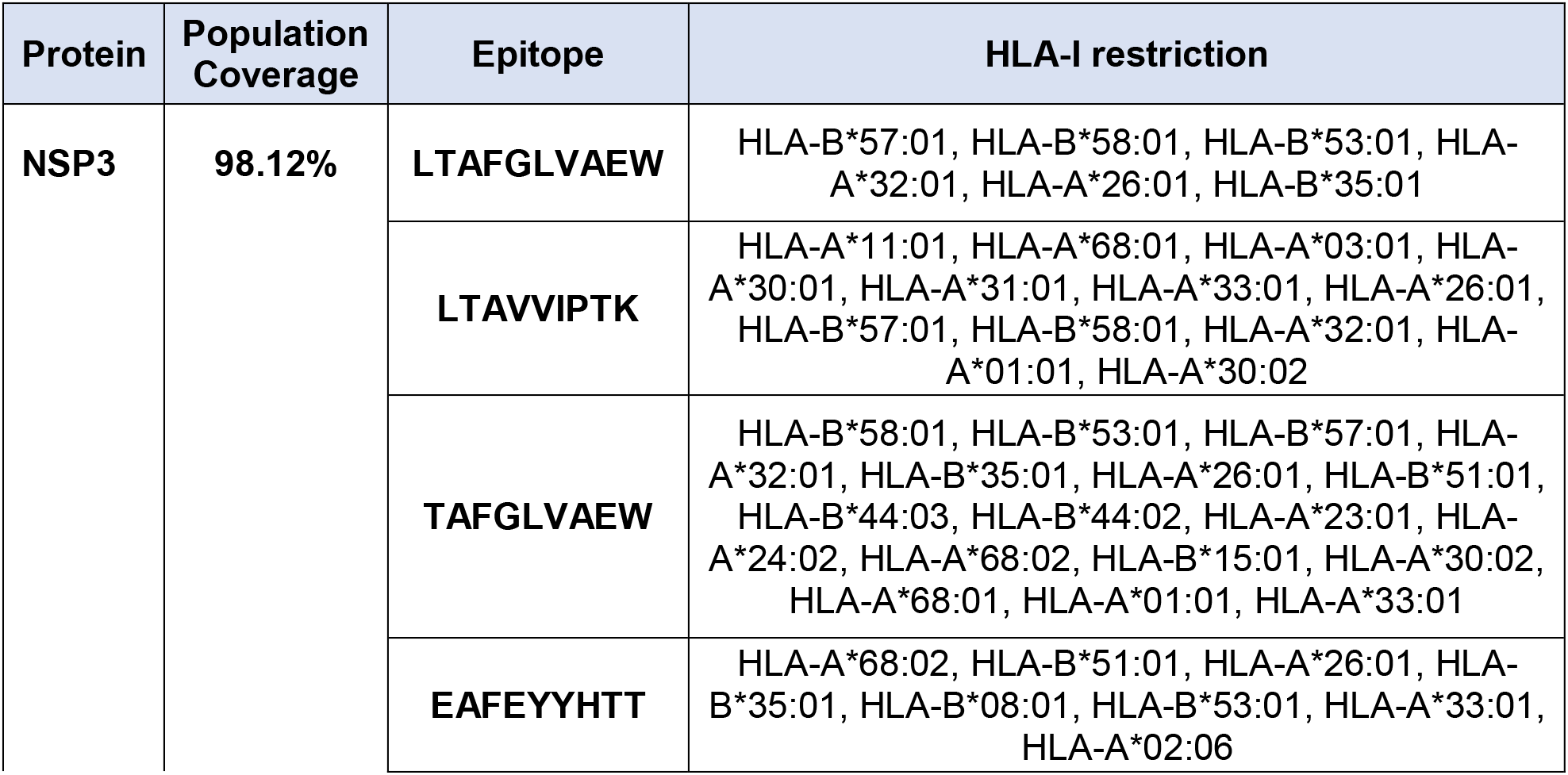
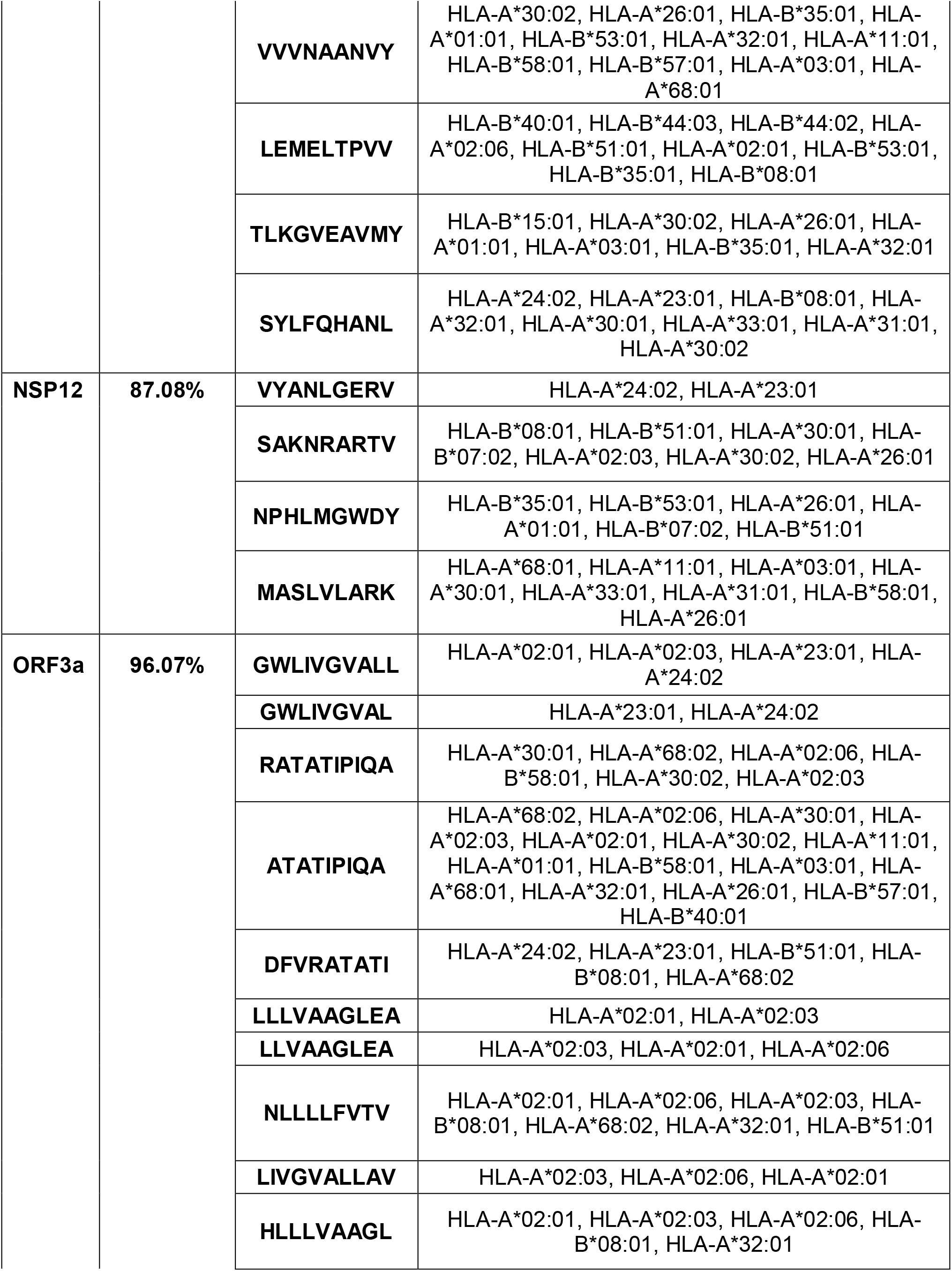
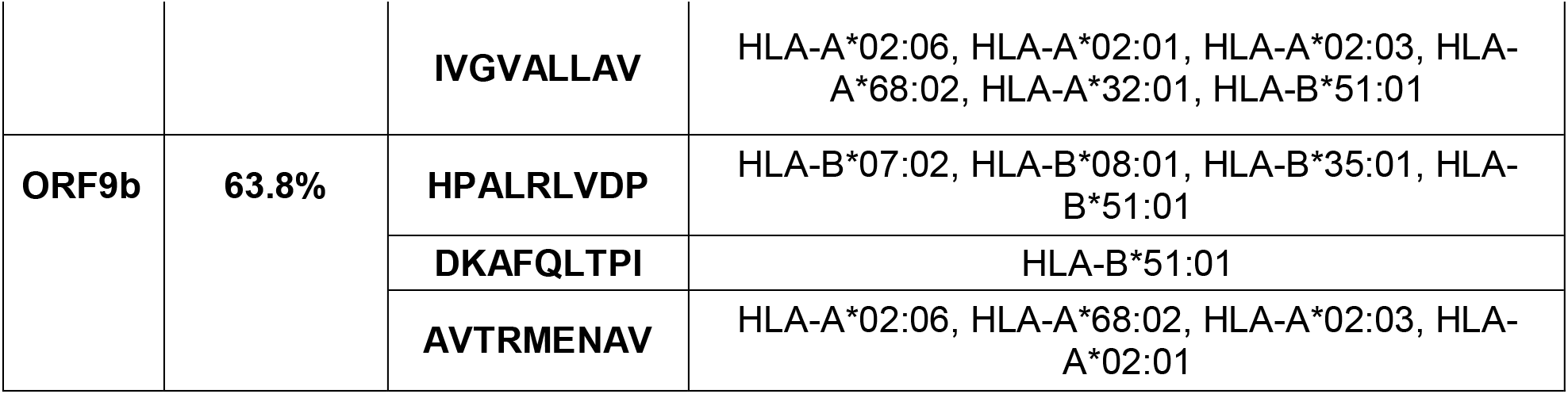
Top common IFN-y inducing CD8+ T cell epitopes from nsp3, nsp12, ORF3a and ORF9b proteins across VOC and their population coverage. The selected epitopes are immunogenic, antigenic, non-allergenic, non-toxic and stable.

| **Protein** | **Population Coverage** | **Epitope** | **HLA-I restriction** |
| --- | --- | --- | --- |
| **NSP3** | **98.12%** | **LTAFLGLVAEW** | HLA-B*57:01, HLA-B*58:01, HLA-B*53:01, HLA-A*32:01, HLA-A*26:01, HLA-B*35:01 |
|  |  | **LTAVVIPTK** | HLA-A*11:01, HLA-A*68:01, HLA-A*03:01, HLA-A*30:01, HLA-A*31:01, HLA-A*33:01, HLA-A*26:01, HLA-B*57:01, HLA-B*58:01, HLA-A*32:01, HLA-A*01:01, HLA-A*30:02 |
|  |  | **TAFGLVAEW** | HLA-B*58:01, HLA-B*53:01, HLA-B*57:01, HLA-A*32:01, HLA-B*35:01, HLA-A*26:01, HLA-B*51:01, HLA-B*44:03, HLA-B*44:02, HLA-A*23:01, HLA-A*24:02, HLA-A*68:02, HLA-B*15:01, HLA-A*30:02, HLA-A*68:01, HLA-A*01:01, HLA-A*33:01 |
|  |  | **EAFFEYHTT** | HLA-A*68:02, HLA-B*51:01, HLA-A*26:01, HLA-B*35:01, HLA-B*08:01, HLA-B*53:01, HLA-A*33:01, HLA-A*02:06 |
|  |  | **VVVNAANVY** | HLA-A*30:02, HLA-A*26:01, HLA-B*35:01, HLA-A*01:01, HLA-B*53:01, HLA-A*32:01, HLA-A*11:01, HLA-B*58:01, HLA-B*57:01, HLA-A*03:01, HLA-A*68:01 |
|  |  | **LEMELTPVV** | HLA-B*40:01, HLA-B*44:03, HLA-B*44:02, HLA-A*02:06, HLA-B*51:01, HLA-A*02:01, HLA-B*53:01, HLA-B*35:01, HLA-B*08:01 |
|  |  | **TLKGVEAVMY** | HLA-B*15:01, HLA-A*30:02, HLA-A*26:01, HLA-A*01:01, HLA-A*03:01, HLA-B*35:01, HLA-A*32:01 |
|  |  | **SYLFQHANL** | HLA-A*24:02, HLA-A*23:01, HLA-B*08:01, HLA-A*32:01, HLA-A*30:01, HLA-A*33:01, HLA-A*31:01, HLA-A*30:02 |
| **NSP12** | **87.08%** | **VYANLGERV** | HLA-A*24:02, HLA-A*23:01 |
|  |  | **SAKNRARTV** | HLA-B*08:01, HLA-B*51:01, HLA-A*30:01, HLA-B*07:02, HLA-A*02:03, HLA-A*30:02, HLA-A*26:01 |
|  |  | **NPHLMGWDY** | HLA-B*35:01, HLA-B*53:01, HLA-A*26:01, HLA-A*01:01, HLA-B*07:02, HLA-B*51:01 |
|  |  | **MASLVLARK** | HLA-A*68:01, HLA-A*11:01, HLA-A*03:01, HLA-A*30:01, HLA-A*33:01, HLA-A*31:01, HLA-B*58:01, HLA-A*26:01 |
| **ORF3a** | **96.07%** | **GWLIVGVALL** | HLA-A*02:01, HLA-A*02:03, HLA-A*23:01, HLA-A*24:02 |
|  |  | **GWLIVGVAL** | HLA-A*23:01, HLA-A*24:02 |
|  |  | **RATATIPIQA** | HLA-A*30:01, HLA-A*68:02, HLA-A*02:06, HLA-B*58:01, HLA-A*30:02, HLA-A*02:03 |
|  |  | **ATATIPIQA** | HLA-A*68:02, HLA-A*02:06, HLA-A*30:01, HLA-A*02:03, HLA-A*02:01, HLA-A*30:02, HLA-A*11:01, HLA-A*01:01, HLA-B*58:01, HLA-A*03:01, HLA-A*68:01, HLA-A*32:01, HLA-A*26:01, HLA-B*57:01, HLA-B*40:01 |
|  |  | **DFVRATATI** | HLA-A*24:02, HLA-A*23:01, HLA-B*51:01, HLA-B*08:01, HLA-A*68:02 |
|  |  | **LLLVAAGLEA** | HLA-A*02:01, HLA-A*02:03 |
|  |  | **LLVAAGLEA** | HLA-A*02:03, HLA-A*02:01, HLA-A*02:06 |
|  |  | **NLLLLFVTV** | HLA-A*02:01, HLA-A*02:06, HLA-A*02:03, HLA-B*08:01, HLA-A*68:02, HLA-A*32:01, HLA-B*51:01 |
|  |  | **LIVGVALLAV** | HLA-A*02:03, HLA-A*02:06, HLA-A*02:01 |
|  |  | **HLLLVAAGL** | HLA-A*02:01, HLA-A*02:03, HLA-A*02:06, HLA-B*08:01, HLA-A*32:01 |
|  |  | **IVGVALLAV** | HLA-A*02:06, HLA-A*02:01, HLA-A*02:03, HLA-A*68:02, HLA-A*32:01, HLA-B*51:01 |
| **ORF9b** | **63.8%** | **HPALRLVDP** | HLA-B*07:02, HLA-B*08:01, HLA-B*35:01, HLA-B*51:01 |
|  |  | **DKAFQLTPI** | HLA-B*51:01 |
|  |  | **AVTRMENAV** | HLA-A*02:06, HLA-A*68:02, HLA-A*02:03, HLA-A*02:01 |

### CD4+ T cell epitope prediction

IEDB recommended 2.22 (rank) was selected as the best performing tool-attribute combination for variant CD4 epitope predictions due to its sensitivity and specificity computed in a previous study [56]. IFNEpitope, VaxiJen 2.0, AllerTop 2.0, ToxinPred and ProtParam, only antigenic, IFN-y inducing, non-allergenic, non-toxic and stable CD4 epitopes were selected. We obtained 28 common nsp3, 19 common nsp12, 11 common ORF3a and 4 common ORF9b epitopes across VOC, 1 nsp3 epitope specific to the UK variant, 3 nsp3 epitopes specific to the Brazilian variant, 5 ORF3a epitopes specific to the South African variant and 3 ORF3a epitopes specific to the Indian variant (**Supplementary Table 7**).

### Murine MHC restriction prediction

As murine models pose a great way to safely test the immunogenicity and efficacy of potential vaccine constructs, we identified top CD8 epitopes that have either strong or weak affinity to murine H2 molecules (**Table 3**). ORF3a was predicted to have 6 common immunogenic, antigenic, non-allergenic, non-toxic and stable CD8 epitopes across VOC with affinity to murine MHC restriction, while nsp3 and nsp12 had 5 and 4 CD8 epitopes, respectively (**Table 3**). Moreover, our analysis revealed three common immunogenic, antigenic, non-toxic, non-allergenic, stable and IFN-y inducing CD8 epitopes presented by murine MHC restriction that overlap with at least one antigenic, non-allergenic, non-toxic, stable and IFN-y inducing CD4 epitope: GWLIVGVAL (ORF3a), TAFGLVAEW (nsp3) and VYANLGERV (nsp12). These CD8 epitopes were found to be within a total of 6 antigenic, non-allergenic, non-toxic, stable and IFN-y inducing CD4 epitopes: TAFGLVAEWFLAYIL (nsp3), LTAFGLVAEWFLAYI (nsp3), DLTAFGLVAEWFLAY (nsp3), PDILRVYANLGERVR (nsp12), RVYANLGERVRQALL (nsp12) and PFGWLIVGVALLAVF (ORF3a) (**Table 4**). The world population coverage analysis revealed that GWLIVGVAL, TAFGLVAEW and VYANLGERV are able to elicit an immune response that covers 75.72% of the world population (**Supplementary Figure 2p**) while the combined 9 HLA-I and HLA-II epitopes are able to cover 99.99% of the world population (**Supplementary Figure 2q**). Therefore, we can state that the presented overlapping CD8 and CD4 epitopes could act as an excellent multi-epitope vaccine construct against SARS-CoV-2.

**Table 3.**
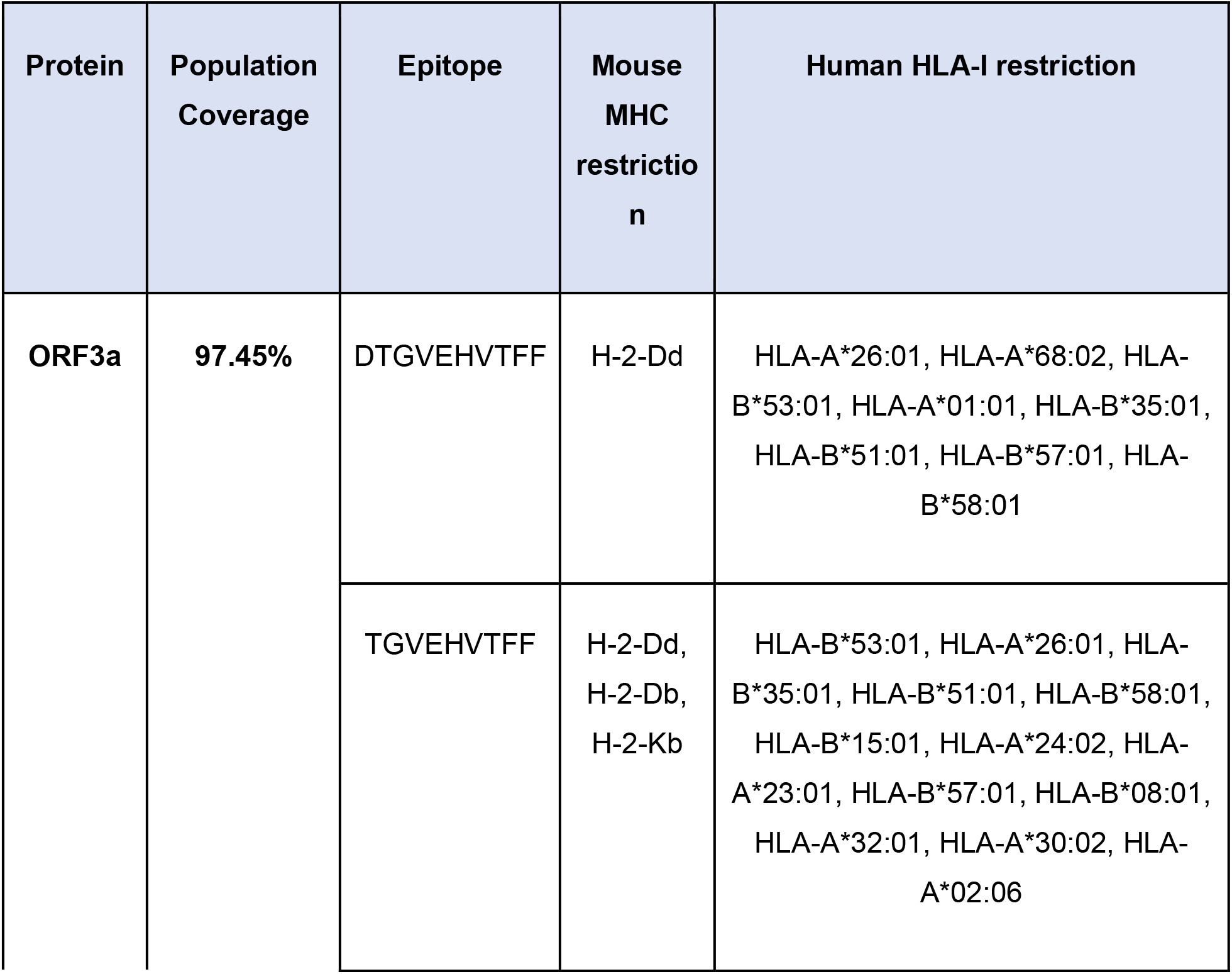
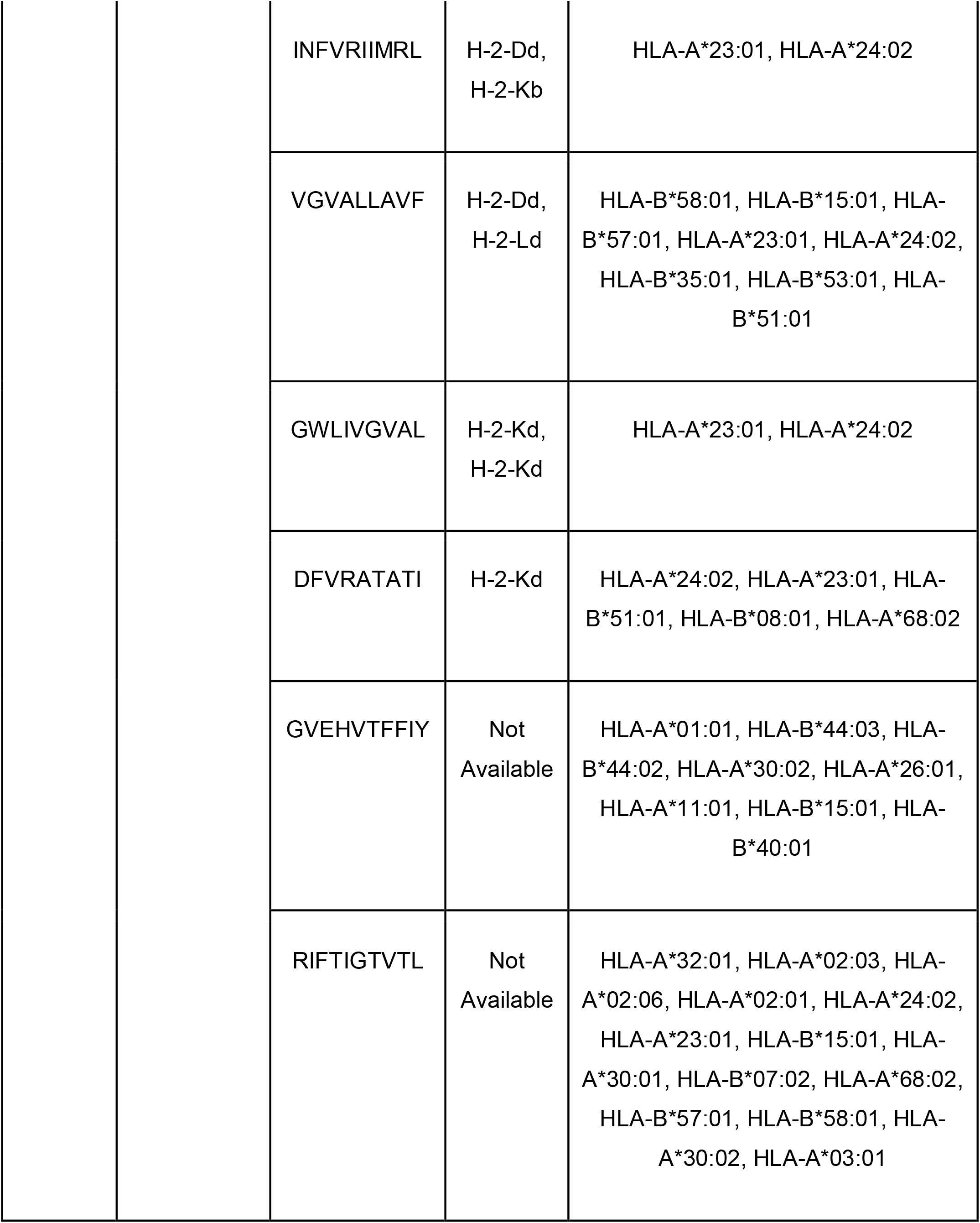
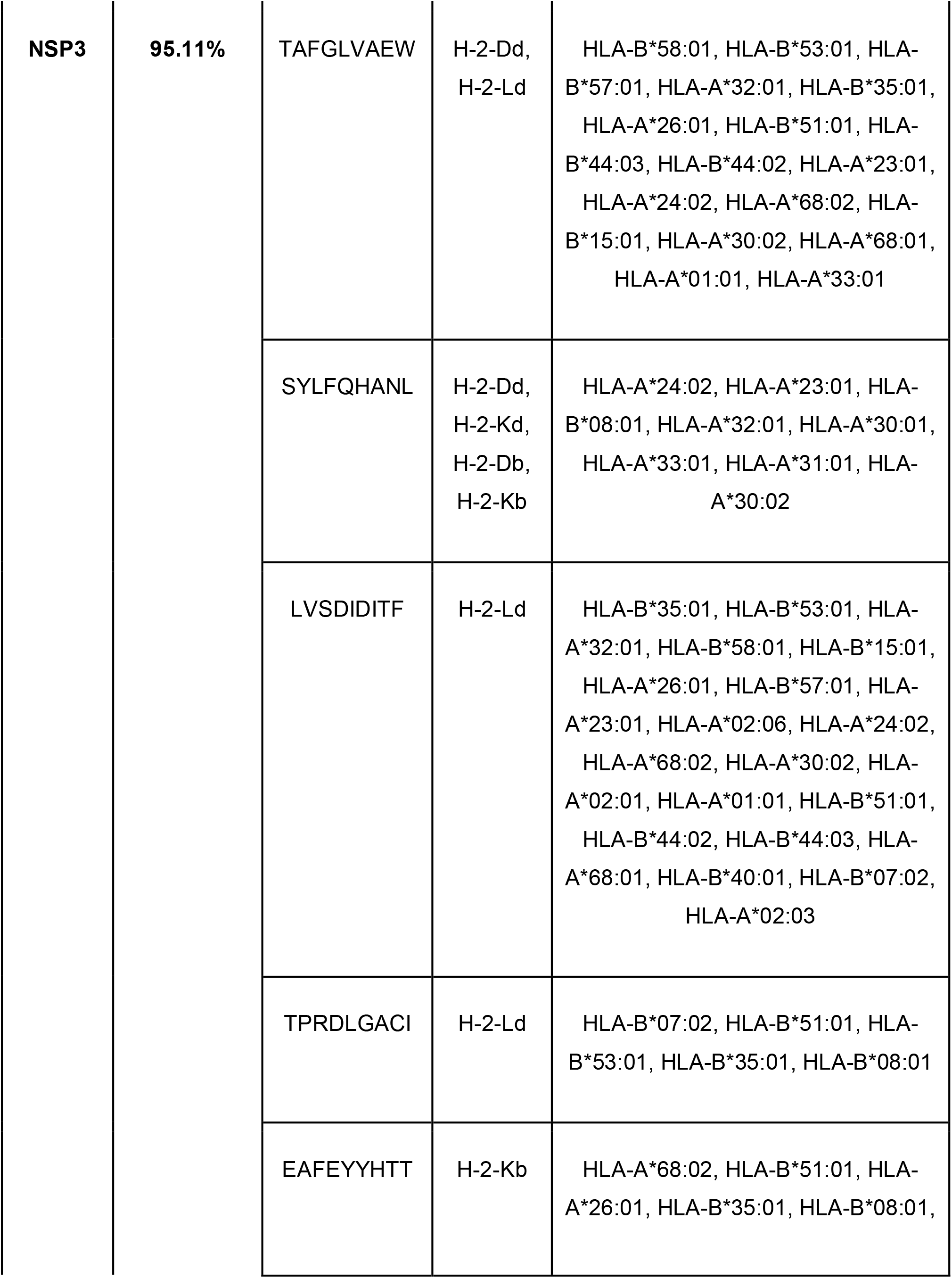
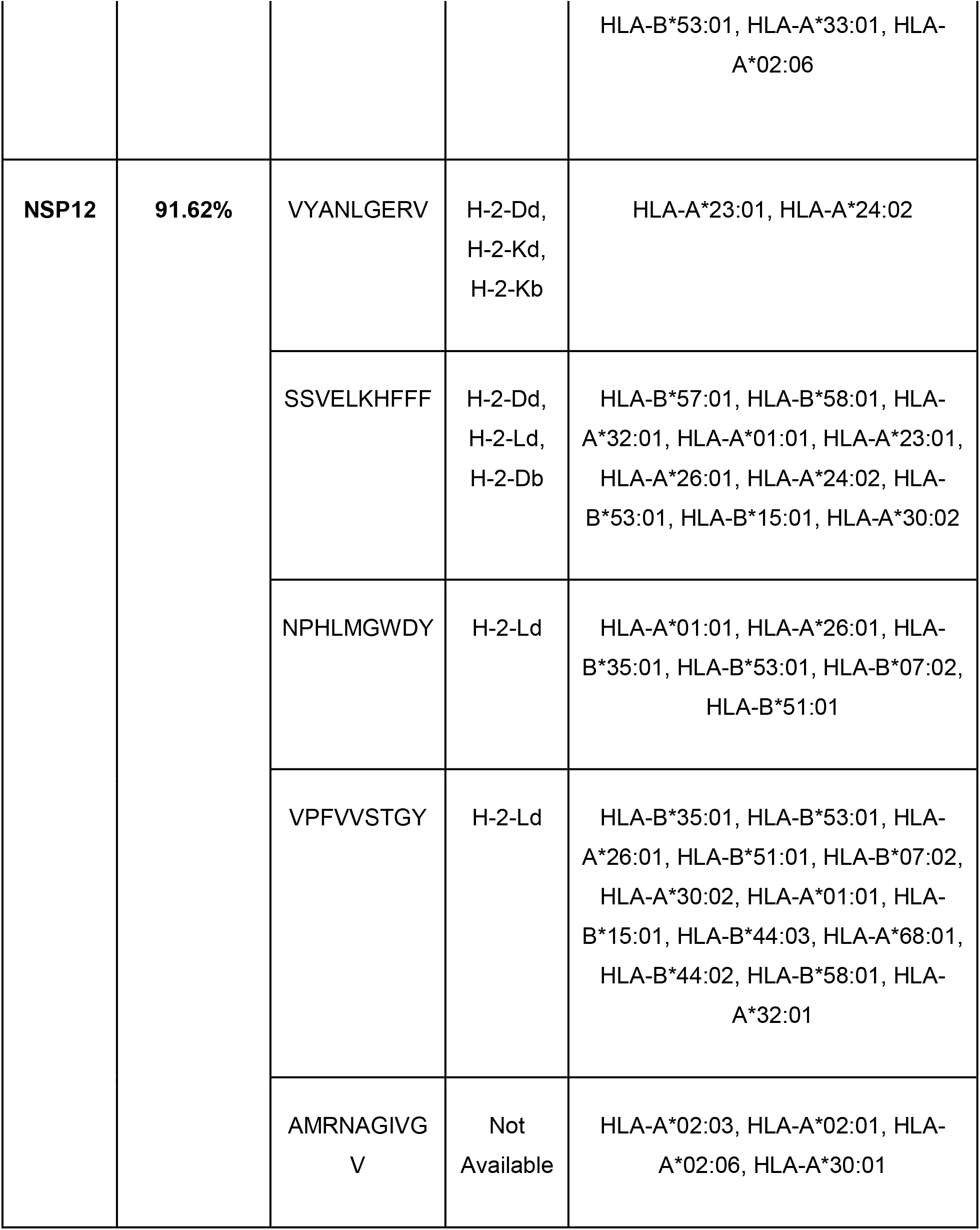
Top common immunogenic nsp3, nsp12 and ORF3a CD8+ T cell epitopes to all four variants of concern in SARS-CoV-2 and their population coverage. These epitopes are antigenic, non-allergenic, non-toxic and stable. Epitopes that are presented on murine MHC were prioritized for selection to population coverage analysis. No ORF9b-derived top epitopes were found to have either a weak or strong binding affinity to a mouse MHC.

**Table 4.** Top common antigenic CD4+ T cell epitopes across VOC and their population coverage. The selected epitopes are IFN-y inducing, non-allergenic, non-toxic and stable.

| Protein | Population Coverage | Epitope | HLA-II restriction |
| --- | --- | --- | --- |
| **ORF3a** | 99.76% | GVHFVCNLLLLFVTV | HLA-DPA1*03:01/DPB1*04:02, HLA-DPA1*02:01/DPB1*01:01, HLA-DPA1*01:03/DPB1*02:01, HLA-DPA1*02:01/DPB1*05:01 |
|  |  | FVTVYSHLLLVAAGL | HLA-DRB1*01:01, HLA-DPA1*03:01/DPB1*04:02, HLA-DQA1*01:02/DQB1*06:02, HLA-DRB1*07:01, HLA-DPA1*01:03/DPB1*04:01, HLA-DRB1*15:01 |
|  |  | VHFVCNLLLLFVTVY | HLA-DPA1*03:01/DPB1*04:02, HLA-DPA1*01:03/DPB1*02:01, HLA-DPA1*02:01/DPB1*01:01, HLA-DPA1*02:01/DPB1*05:01 |
|  |  | HLLLVAAGLEAPFLY | HLA-DQA1*05:01/DQB1*02:01 |
|  |  | SHLLLVAAGLEAPFL | HLA-DQA1*05:01/DQB1*02:01 |
|  |  | PFGWLIVGVALLAVF | HLA-DPA1*03:01/DPB1*04:02, HLA-DRB1*01:01, HLA-DQA1*05:01/DQB1*02:01, HLA-DRB1*08:02, HLA-DQA1*01:02/DQB1*06:02, HLA-DQA1*05:01/DQB1*03:01 |
|  |  | LLLVAAGLEAPFLYL | HLA-DQA1*05:01/DQB1*02:01, HLA-DQA1*05:01/DQB1*03:01 |
| **NSP3** | 99.82% | LTAFGGLVAEWFLAYI | HLA-DPA1*01:03, HLA-DPB1*02:01, HLA-DQA1*05:01, HLA-DQB1*02:01, HLA-DPA1*01:03, HLA-DPB1*04:01, HLA-DQA1*01:01, HLA-DQB1*05:01, HLA-DPA1*03:01, HLA-DPB1*04:02, HLA-DPA1*02:01, HLA-DPB1*01:01, HLA-DQA1*03:01, HLA-DQB1*03:02 |
|  |  | PTVVVNAAENVYLKHG | HLA-DRB1*13:02, HLA-DRB1*08:02, HLA-DQA1*01:02, HLA-DQB1*06:02 |
|  |  | DLTAFGLVAEWFLAY | HLA-DPA1*01:03, HLA-DPB1*02:01, HLA-DQA1*05:01, HLA-DQB1*02:01, HLA-DPA1*01:03, HLA-DPB1*04:01, HLA-DPA1*03:01, HLA-DPB1*04:02, HLA-DQA1*03:01, HLA-DQB1*03:02, HLA-DQA1*04:01, HLA-DQB1*04:02, HLA-DQA1*01:01, HLA-DQB1*05:01, HLA-DPA1*02:01, HLA-DPB1*01:01 |
| **NSP12** | 96.62% | NLKYAISAKNRARTV | HLA-DPA1*02:01, HLA-DPB1*14:01, HLA-DRB1*11:01, HLA-DRB1*09:01 |
|  |  | LKYAISAKNRARTVA | HLA-DPA1*02:01, HLA-DPB1*14:01, HLA-DRB1*11:01 |
|  |  | PDILRVYANLGERVR | HLA-DRB1*15:01, HLA-DRB1*13:02, HLA-DRB1*04:05 |
|  |  | RVYANLGERVRQALL | HLA-DRB3*02:02 |
|  |  | GLVASIKNFKSVLYY | HLA-DRB1*15:01, HLA-DRB1*12:01, HLA-DRB1*04:01, HLA-DRB1*08:02 |
|  |  | VASIKNFKSVLYYQN | HLA-DRB1*15:01, HLA-DRB1*12:01, HLA-DPA1*02:01, HLA-DPB1*05:01, HLA-DRB1*08:02, HLA-DPA1*01:03, HLA-DPB1*04:01, HLA-DRB1*04:01, HLA-DRB1*04:05 |
| **ORF9b** | 94.74% | KAFQLTPIAVQMTKL | HLA-DPA1*02:01, HLA-DPB1*14:01, HLA-DRB4*01:01, HLA-DRB1*04:01, HLA-DRB1*01:01 |
|  |  | LEDKAFQLTPIAVQM | HLA-DPA1*02:01, HLA-DPB1*14:01, HLA-DPA1*01:03, HLA-DPB1*04:01 |
|  |  | SLEDKAFQLTPIAVQ | HLA-DPA1*02:01, HLA-DPB1*14:01 |
|  |  | NSLEDKAFQLTPIAV | HLA-DQA1*01:01, HLA-DQB1*05:01 |

### Population Coverage Analysis

#### CD8 epitopes

The population coverage of the T cell epitopes was estimated from the corresponding HLA associations. The population coverage was calculated for both datasets with and without IFNEpitope filter. The top common 8 nsp3, 4 nsp12, 11 ORF3a and 3 ORF9b epitopes across VOC cover 98.12%, 87.08%, 96.07% and 63.8% of the world population, respectively (**Table 2, Supplementary Figures 2e-h)**, whereas all of them are immunogenic, antigenic, non-allergenic, non-toxic, stable and IFN-y inducing. Since IFNEpitope is more accurate for CD4 epitope predictions and in order to increase the population coverage, IFN-y secretion filter was removed from the predictions. It was found that the 21 nsp3, 19 nsp12, 36 ORF3a and 5 ORF9b are able to elicit an immune response that covers 98.55% of world population in nsp3, nsp12 and ORF3a proteins, and 75.8% in ORF9b (**Supplementary Table 2, Supplementary Figures 2a-d)**. Als, we computed the population coverage for our top common CD8 epitopes that are presented by murine model MHC restriction (**Table 3**). The fewer epitope-construct is still able to elicit a strong immune response that covers 97.45% of population in ORF3a, 95.11% in nsp3 and 91.62% in nsp12 world population (**Table 3, Supplementary Figures 2i-k**).

#### CD4 epitopes

The population coverage of CD4 epitopes was calculated for the few top epitopes. It was found that the top common 7 ORF3a, 3 nsp3, 6 nsp12 and 4 ORF9b epitopes are able to cover 99.76%, 99.82%, 96.62% and 94.74% of the world population, respectively (**Table 4, Supplementary Figures 2l-o)**.

#### 3D structure predictions

The top CD8 epitopes from each protein were selected and visualized with a respective HLA-I allele (**Figures 5-8**). TAFGLAEW was chosen as the best epitope in nsp3 due to its high antigenicity score of 1.0431, immunogenicity score of 0.19168, stability score of -7.81 (**Supplementary Table 3**) and its presence in two antigenic CD4 epitopes identified in this study (**Table 5**). For nsp12, SSVELKHFFF was chosen due to its high antigenicity score of 1.4503 (**Supplementary Table 5**). GWLIVGVAL was selected as the best CD8 epitope in ORF3a due to its high immunogenicity score of 0.26012, positive IFN-y score (**Supplementary Table 4**) and presence in an antigenic CD4 epitope (**Table 4**). Finally, a CD8 epitope DKAFQLTPI from ORF9b was chosen for 3D visualization due to its ability to interact with TOM70. Besides this, DKAFQLTPI was incorporated into two antigenic CD4 epitopes which suggests its higher potential to elicit a protective and strong immune response (**Table 4**). We also visualized the positions of selected top common immunogenic, antigenic, non-allergenic, non-toxic and stable CD8 epitopes across VOC in nsp3, nsp12 and ORF3a reference protein sequences (**Supplementary Figures 1a-c)**. For nsp3, 5CD8 epitopes were visualized: LTAFGLVAEW, TAFGLVAEW, SYLFQHANL, EAFEYYHTT and LVSDIDITF (**Supplementary Figure 1b)**. For nsp12, VYANLGERV, NPHLMGWDY, SSVELKHFFF and SAKNRARTV were visualized (**Supplementary Figure 1c**), and LLVAGLEA, GWLIVGVAL, GWLIVGVALL, NLLLLFVTV, GVEHVTFFIY, VEHVTFFIY, DTGVEHVTFF, GVEHVTFFI and TGVEHVTFF were visualized in ORF3a protein (**Supplementary Figure 1a**). The highest number CD8 epitopes from ORF3a protein was selected for visualization due to its highest number of common immunogenic CD8 epitopes.

**Table 5.**
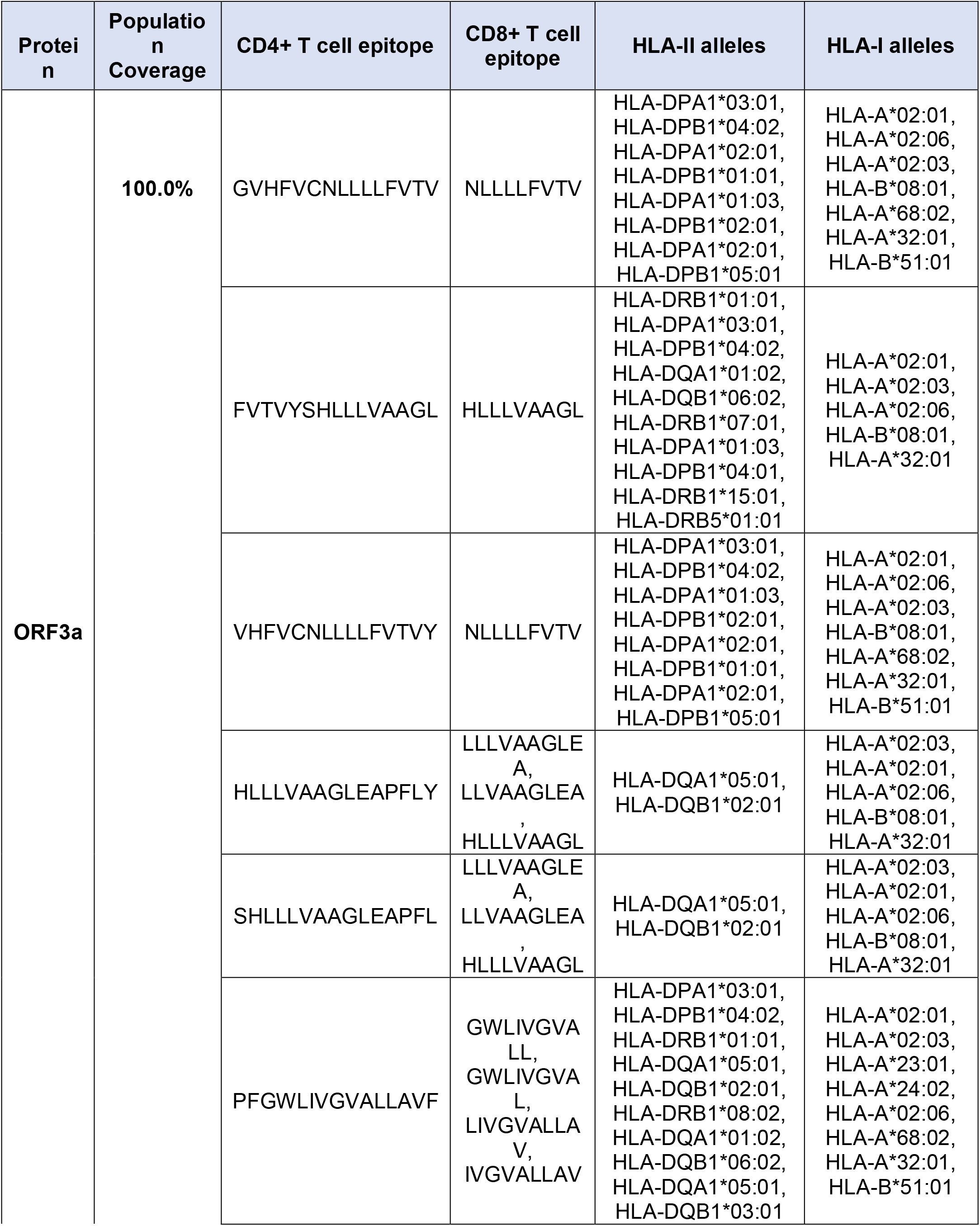
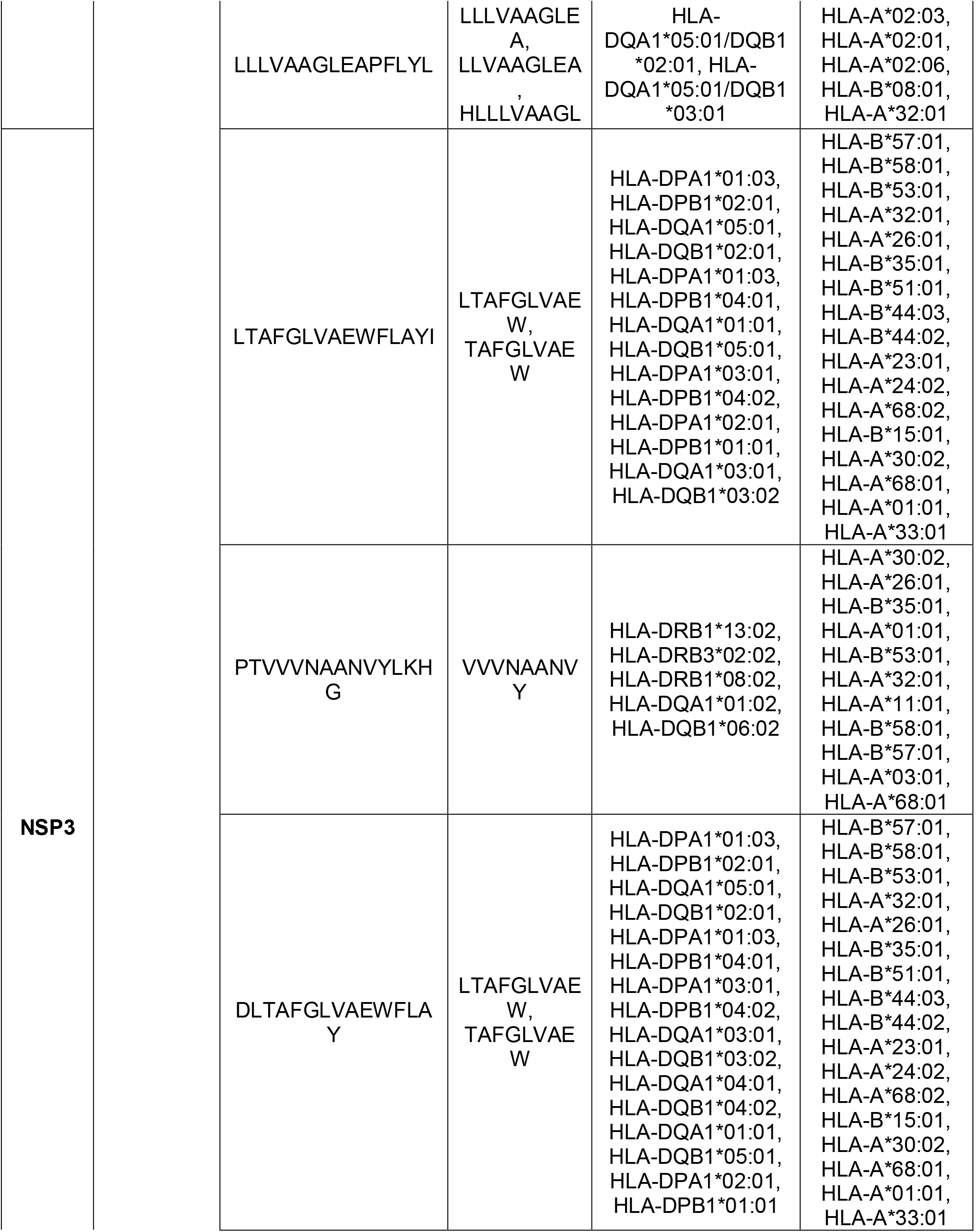
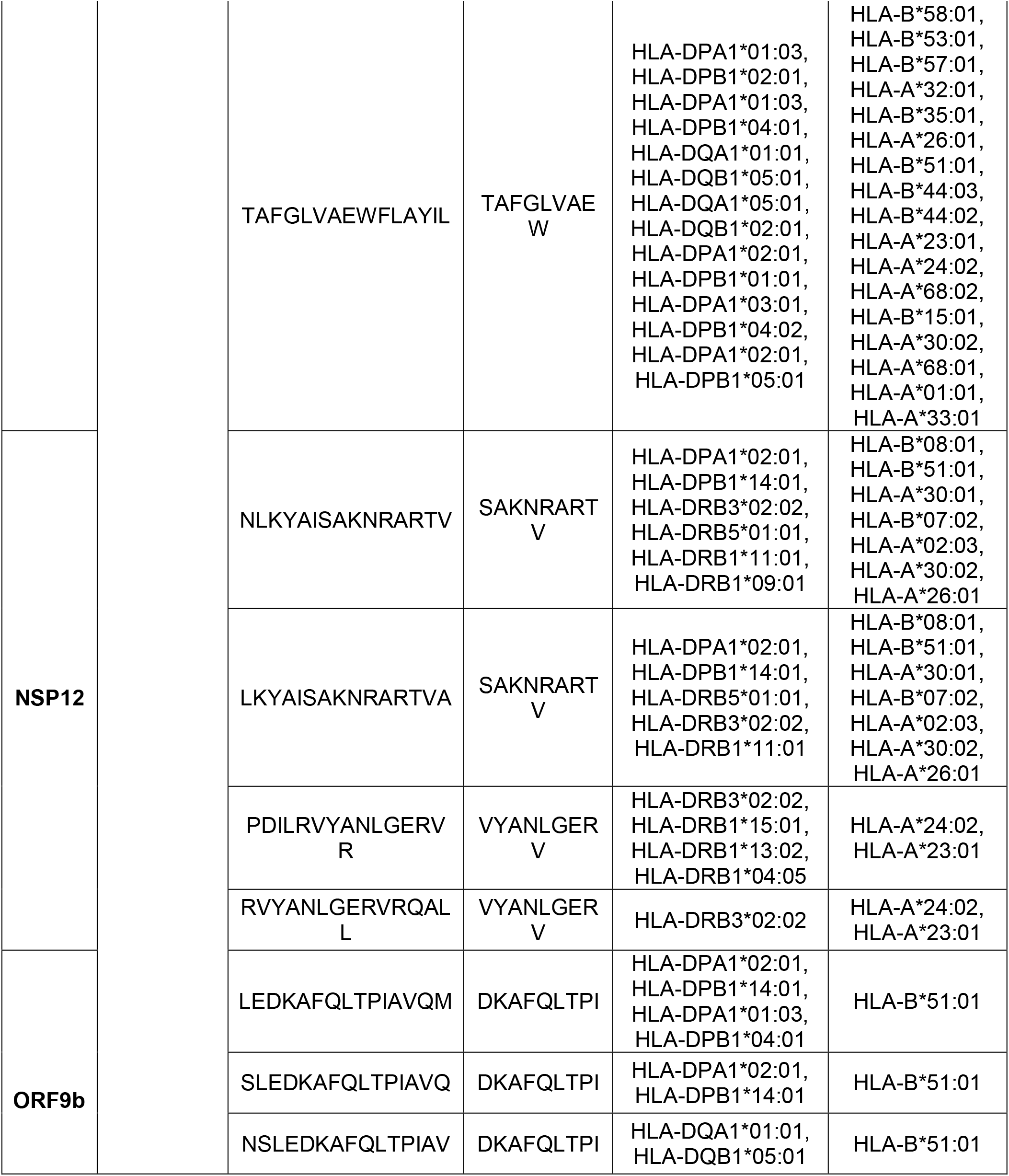
Top overlapping immunogenic, antigenic, non-allergenic, non-toxic, stable, IFN-y inducing CD8 epitopes and antigenic, non-allergenic, non-toxic, stable, IFN-y inducing CD4 epitopes, common across VOC. Population coverage combines both MHC class I and II presented epitopes across nsp3, nsp12, ORF3a and ORF9b.

**Figure 5.**
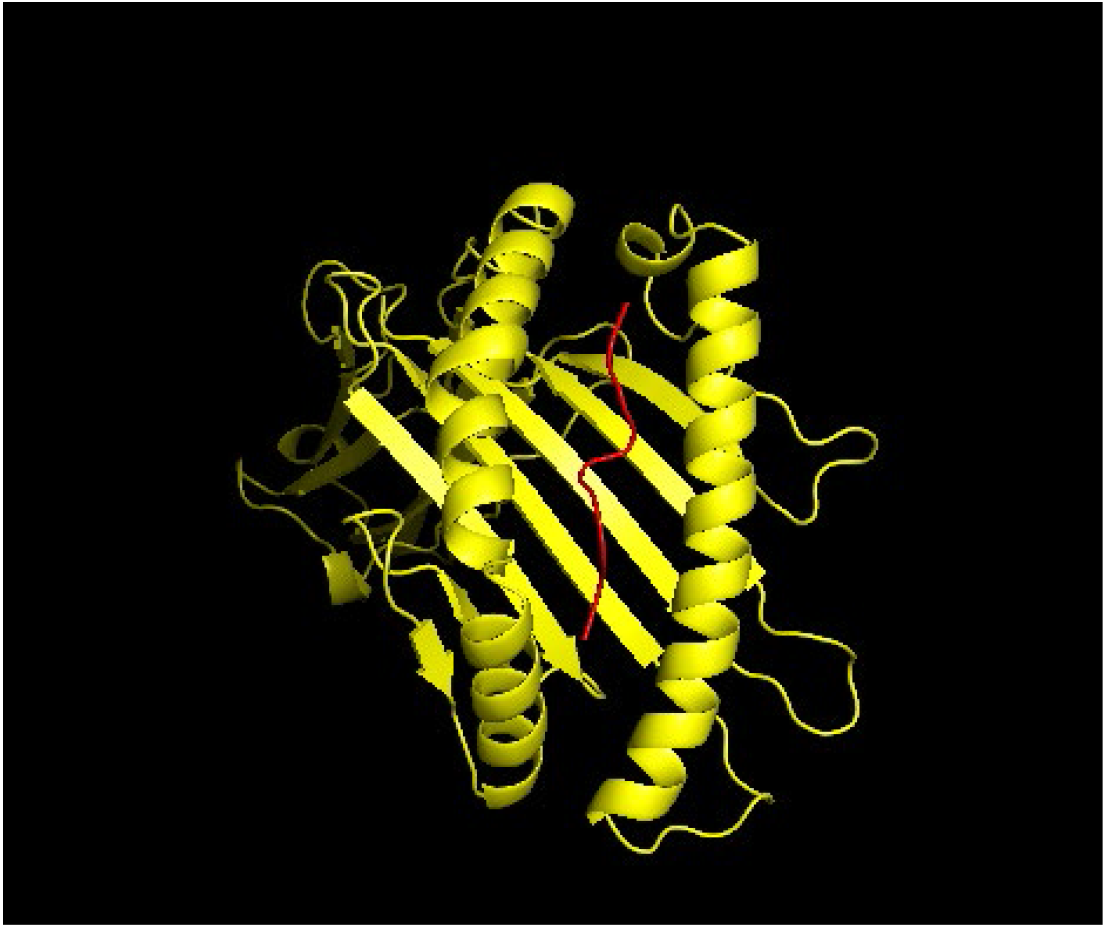
3D structure of TAFGLAEW (red) in nsp3 protein and its interaction with HLA-B*58:01 allele (yellow)

**Figure 6.**
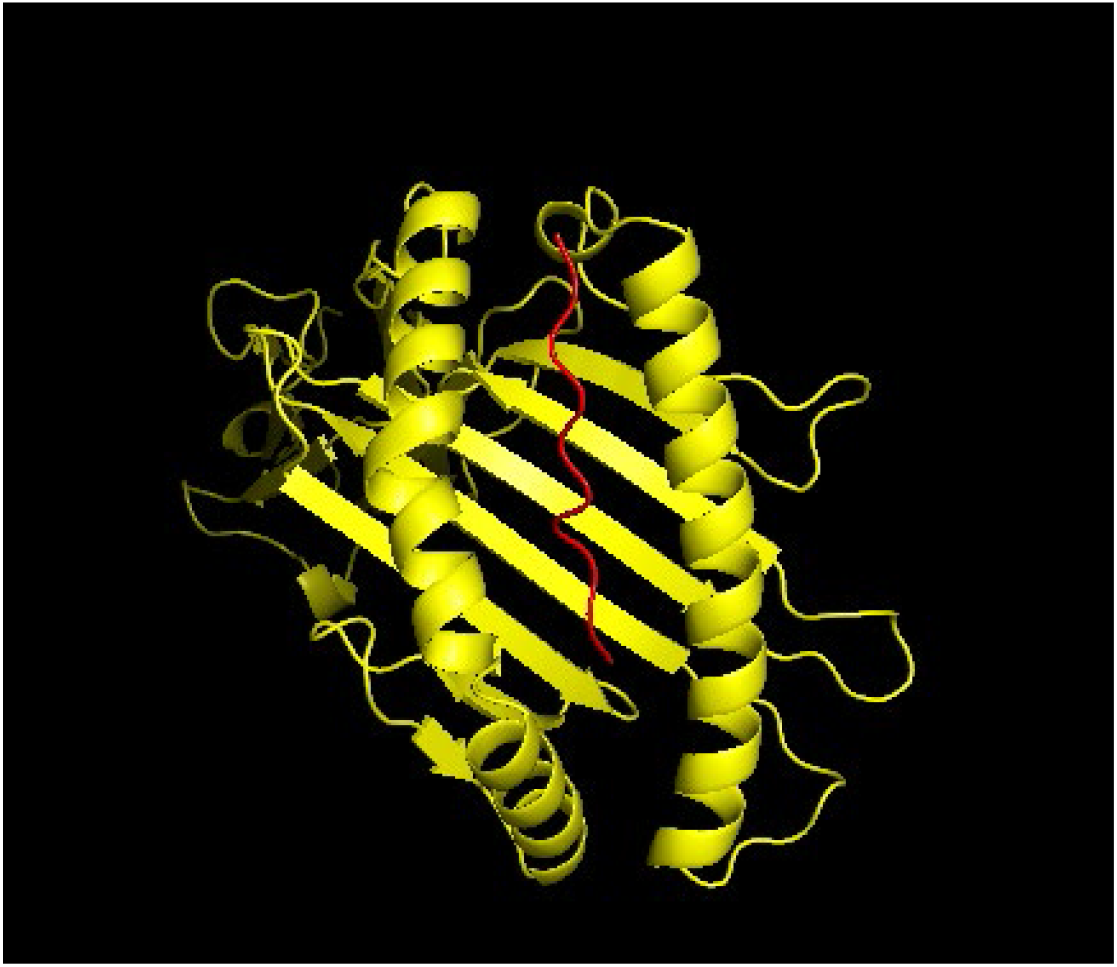
3D structure of SSVELKHFFF (red) in nsp12 protein and its interaction with HLA-B*57:01 allele (yellow)

**Figure 7.**
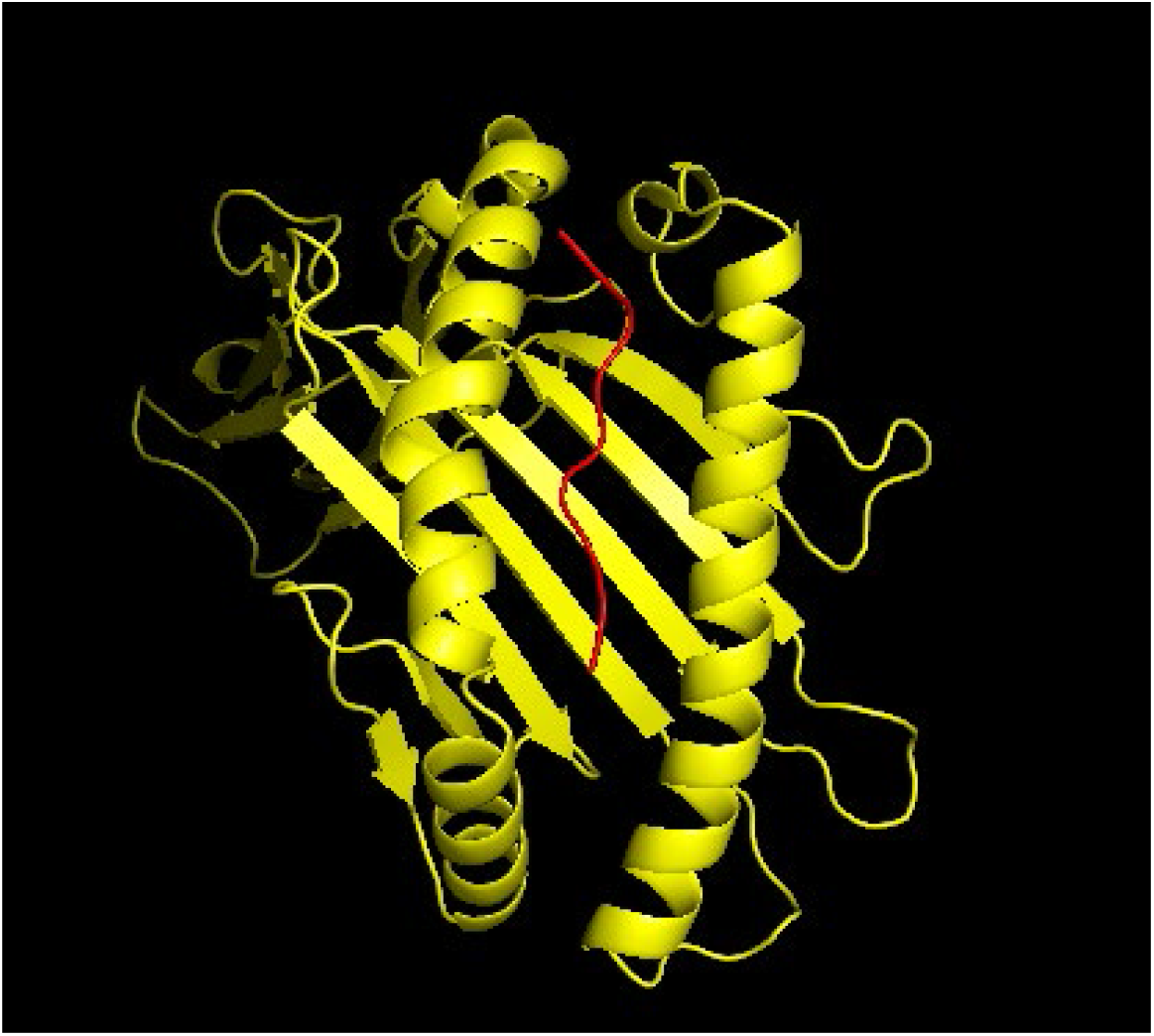
3D structure of GWLIVGVAL (red) in ORF3a protein and its interaction with HLA-A*24:02 allele (yellow)

**Figure 8.**
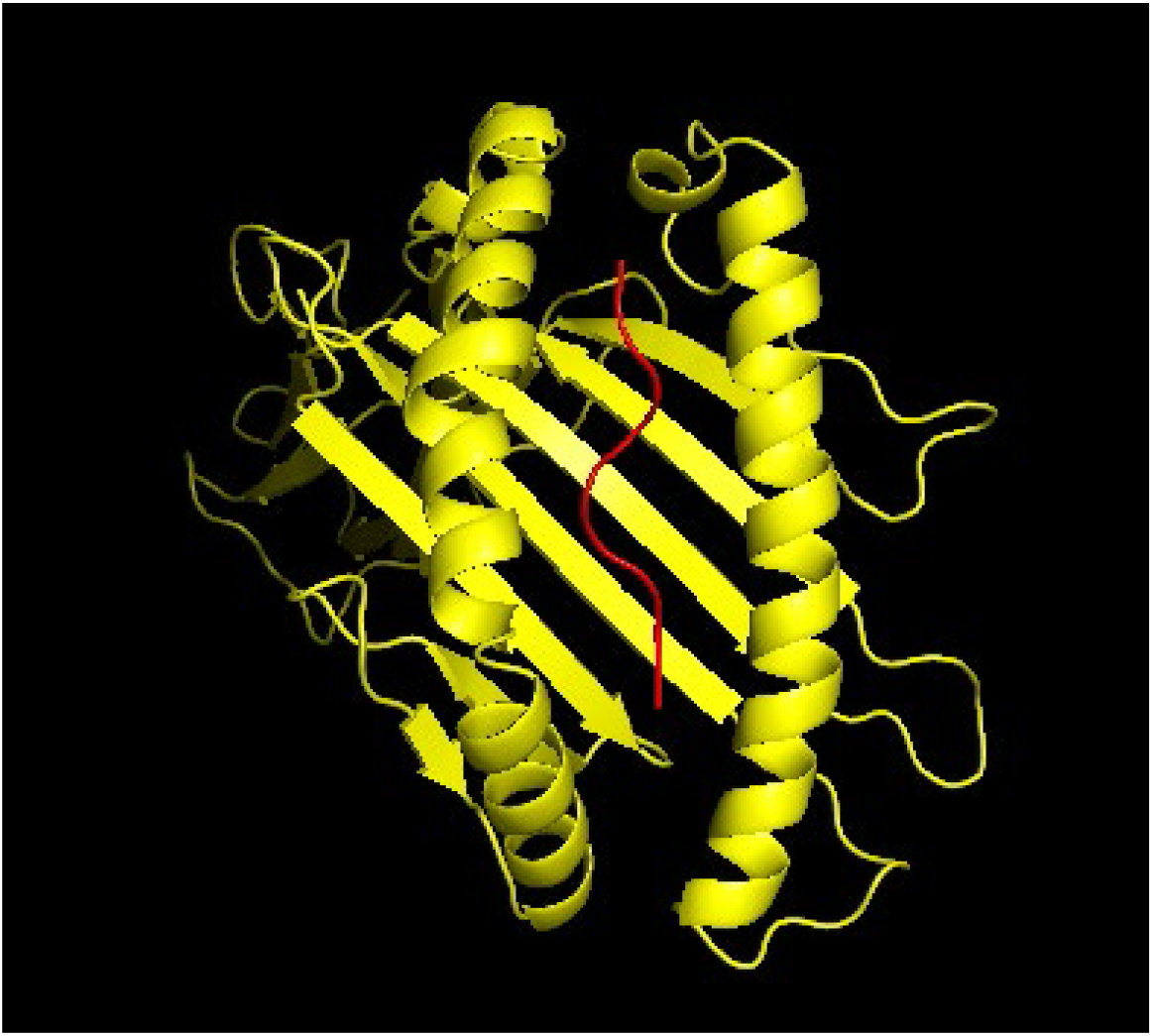
3D structure of DKAFQLTPI (red) in ORF9b protein and its interaction with HLA-B*51:01 allele (yellow)

## Discussion

Built on the basis of immunoinformatics, peptide-based vaccine design has recently become a novel approach towards the identification and design of immunogenic peptides [60]. It is becoming increasingly apparent that T cells, one of the main aspects of the adaptive immune system, may play a significant role in providing protective immunity against COVID-19. Understanding the repertoire of SARS-CoV-2-specific epitopes that can trigger a robust T cell response in humans, is highly important for protective and effective vaccine design (**Figure 1**). With most of the vaccine design attempts focusing on the structural proteins, notably the spike protein, our work, aims to identify immunogenic epitopes from previously uncharacterized ORF proteins. ORF1ab polyprotein, located on the 5’ end of the SARS-CoV-2 genome (**Figure 2**), has been reported to be immunogenic due to its high allele capacity, as well as for its large gene region, compromising of multiple non-structural proteins [7]. PLpro and RdRP, two non-structural proteins from ORF1ab, have essential functions to the virus as they play an important role in viral replication. In addition to these, ORF3a protein, that induces apoptosis in transfected and infected cells, has also been recognized to have high immunogenicity in activating SARS-CoV-2-specific T cells. Choosing the appropriate epitopes that would be safe but also trigger an immune response against an invading pathogen, is a critical step in the development of a multi-epitope-based vaccine. To ensure the reliability of the various prediction tools offered online, tool comparison was the first necessary step towards the identification of the best performing tool-attribute combination to predict T cell epitopes. Our analysis which relied on experimental epitopes matched to the predicted epitopes, revealed that NetMHCpan EL 4.1 ordered based on rank was the best performing tool for CD8 epitope predictions, whereas IEDB recommended 2.22 ordered based on rank was found to perform best for CD4 epitope predictions.

In the present study, we have identified several immunogenic, non-allergenic, non-toxic, stable and IFN-y inducing CD8+ T cell epitopes and antigenic, non-allergenic, non-toxic, stable and IFN-y inducing CD4+ T cell epitopes from SARS-CoV-2 proteins. NSP3 which acts as papain-like protease protein, is responsible for the cleavage of several other non-structural proteins in SARS-CoV-2, an indication of its importance in the virus life cycle [61]. Presence of a macro domain in NSP3 which is involved in disruption of the expression of innate immunity genes, is another indication of the importance of the protein [62]. Our analysis revealed a high number of T cell epitopes in NSP3 with a total of 8 CD8 (**Supplementary Table 3**) and 28 CD4+ T cell epitopes, common to the VOC (**Supplementary Table 7**). The predicted CD8 epitopes had a high presentation of HLA alleles which may contribute to induced immunogenicity [63]. Moreover, we identified 4 antigenic, non-allergenic, non-toxic, stable and IFN-y inducing CD4 epitopes that had a corresponding immunogenic, non-allergenic, non-toxic, stable and IFN-y inducing CD8+ T cell epitope within it (**Table 5**). The combination of CD4 and CD8 T cell epitopes indicates a potential more powerful immune response against SARS-CoV-2.

ORF3a protein, known for the suppression of innate immune response [64], was also predicted to have a high number of T cell epitopes for an effective vaccine design. We predicted a total of 11 immunogenic, non-allergenic, non-toxic, stable and IFN-y inducing CD8 and 11 antigenic, non-allergenic, non-toxic, stable and IFN-y inducing CD4 epitopes in ORF3a protein, common across VOC (**Supplementary Tables 4**,**7**). ORF3a top common epitopes were also predicted to have a wide range of HLA-I and HLA-II binding capacity. Out of the 11 CD4+ T cell epitopes, 7 of them had an immunogenic CD8+ T cell within its sequence (**Table 5**) which indicates these epitopes from ORF3a have a high potential of triggering a strong protective immune response against SARS-CoV-2. Additionally, the population coverage of the 7 IFN-y inducing CD4+ T epitopes was predicted to be 99.76% of the world population (**Table 4**), making ORF3a a desirable vaccine candidate. The results suggested that NSP12 had the lowest number of immunogenic CD8+ T cell epitopes with the lowest population coverage, mainly due to its shorter sequence (932 aa) compared to NSP3 (**Table 2**). However, as CD8 epitope predictions were conducted without the IFN-y server, we were able to obtain 15 additional common immunogenic, antigenic, non-allergenic, non-toxic and stable epitopes (**Supplementary Table 5**) which subsequently boosted the population coverage to 98.55% (**Supplementary Table 2**).

ORF9b is an important source of T cell epitopes that remains largely unexplored in the context of T-cell immunity. Although relatively short (97 aa), ORF9b yielded three immunogenic, non-allergenic, non-toxic, stable and IFN-y inducing HLA-I peptides: HPALRLVDP, DKAFQLTPI and AVTRMENAV that cover 63.8% of the world population (**Supplementary Table 6**). To make the outcome more specific, we aimed to identify immunogenic epitopes from ORF9b that could bind to TOM70. Previous studies report that ORF9b occupies the hydrophobic pocket on the TOM70 C-terminal domain (CTD), suppressing the IFN-I production. ORF9b is also expressed early in infection which points to better HLA-I presentation and immunogenicity [10]. In the study, we were able to identify one ORF9b CD8+ T cell epitope (DKAFQLTPI) in the range of the interaction site (aa 43-78) with TOM70 (PDB:7DHG; **Supplementary Figure 1d**). DKAFQLTPI that binds only to HLA-B*51:01, was shown to be immunogenic, highly antigenic, non-allergenic, non-toxic, IFN-y inducing and stable. Besides this, we identified 3 antigenic, non-allergenic, non-toxic, stable and IFN-y inducing CD4+ T cell epitopes – LEDKAFQLTPIAVQM, SLEDKAFQLTPIAVQ and NSLEDKAFQLTPIAV that had DKAFQLTPI incorporated into the sequence (**Table 5**). Although, DKAFQLTPI has no weak or strong binding affinity to murine MHC restriction, it could be used as a part of the multi-epitope vaccine. The high quality CD8+ T cell epitope with booster CD4 epitopes could provide a strong immune response early in infection and inhibit the virus from evading. Considering the critical role of IFN-I in the human antiviral response, re-creation of IFN-I production in COVID-19 infected individuals may prove to be a significantly effective therapeutic option. Since ORF9b is also highly homologous to ORF9b protein in SARS-CoV, the vaccine construct could be used against other SARS-like coronaviruses [10].

A limitation of current studies relates to mutational spectra of SARS-CoV-2 that is constantly changing. This poses a risk to the vaccine candidates to no longer being viable to offer protection against newly formed variants. To overcome this, we selected the most prevalent variants of concern and their most common mutations in nsp3, nsp12, ORF3a and ORF9b proteins across all identified sequences to date. The selected proteins may carry some important mutations that increase the transmissibility and survival of the virus, like the nonsynonymous mutation P323L in nsp12 which has possibly affected the catalytic activity of the nsp12 protein, therefore altering the binding capability with other cofactors, such as nsp7 and nsp8 proteins [65]. However, studies have reported generally low mutation rates in these proteins which increase the efficiency of the proposed multi-epitope vaccine candidates in the long term [66]. Another challenge with peptide vaccines is that peptides can be easily degraded by peptidases which causes a lower T cell response. The present study addresses this issue by calculating the half-life of the top common epitopes to make sure that the vaccine construct is not degraded early in blood.

## Conclusion

With the lack of effective antiviral treatments, there is an urgent need for a protective vaccine that is effective against different strains of SARS-CoV-2 and among different ethnic groups across the world. Our work sheds light on previously uncharacterized SARS-CoV-2 HLA-I and HLA-II peptides from ORF proteins in the SARS-CoV-2 genome which reveals multiple T cell epitopes that could be used in peptide-based vaccine construction. To ensure the vaccine construct is effective against most common circulating mutations, most prevalent mutations from four lineages of variants of concern: B.1.1.7, P.1, B.1.617.2 and B.1.351 were collected and analyzed. We predicted CD4 and CD8 T cell epitopes for two non-structural proteins, nsp3 and nsp12, as well as from ORF3a and ORF9b proteins. We identified 5 immunogenic, antigenic, non-allergenic, non-toxic, stable and IFN-y inducing nsp3 CD8 epitopes with at least weak affinity to one or more mouse MHC alleles, 4 for nsp12 protein and 6 for ORF3a protein, all common to the studied VOC. Besides this, the variant specific epitopes could be used in variant-specific booster vaccine constructs. Our analysis also revealed 3 immunogenic, antigenic, non-allergenic, non-toxic, stable and IFN-y inducing CD8 epitopes with affinity to mouse MHC alleles and presence within at least one antigenic non-allergenic, non-toxic, stable and IFN-y inducing CD4 epitope which altogether have the ability to provoke a strong immune response and protect 99.99% of the world population, suggesting its efficiency as potential multi-epitope vaccine construct. ORF9b, the open-reading frame protein within nucleocapsid protein, was predicted to have one immunogenic, antigenic, non-allergenic, non-toxic, stable and IFN-y inducing CD8 epitope, DKAFQLTPI, that could suppress the interaction between ORF9b and human mitochondrial outer membrane protein, TOM70. These findings suggest that a single multi-epitope vaccine candidate should be efficacious against currently circulating lineages.

## Acknowledgements

We wish to acknowledge for the support by Lombardi Comprehensive Cancer Center, Georgetown University Medical Center.

