## Supplementary Figure 1 for "Design of Peptide Vaccine for COVID19: CD8+ and CD4+ T cell epitopes from SARS-CoV-2 open-reading-frame protein variants"

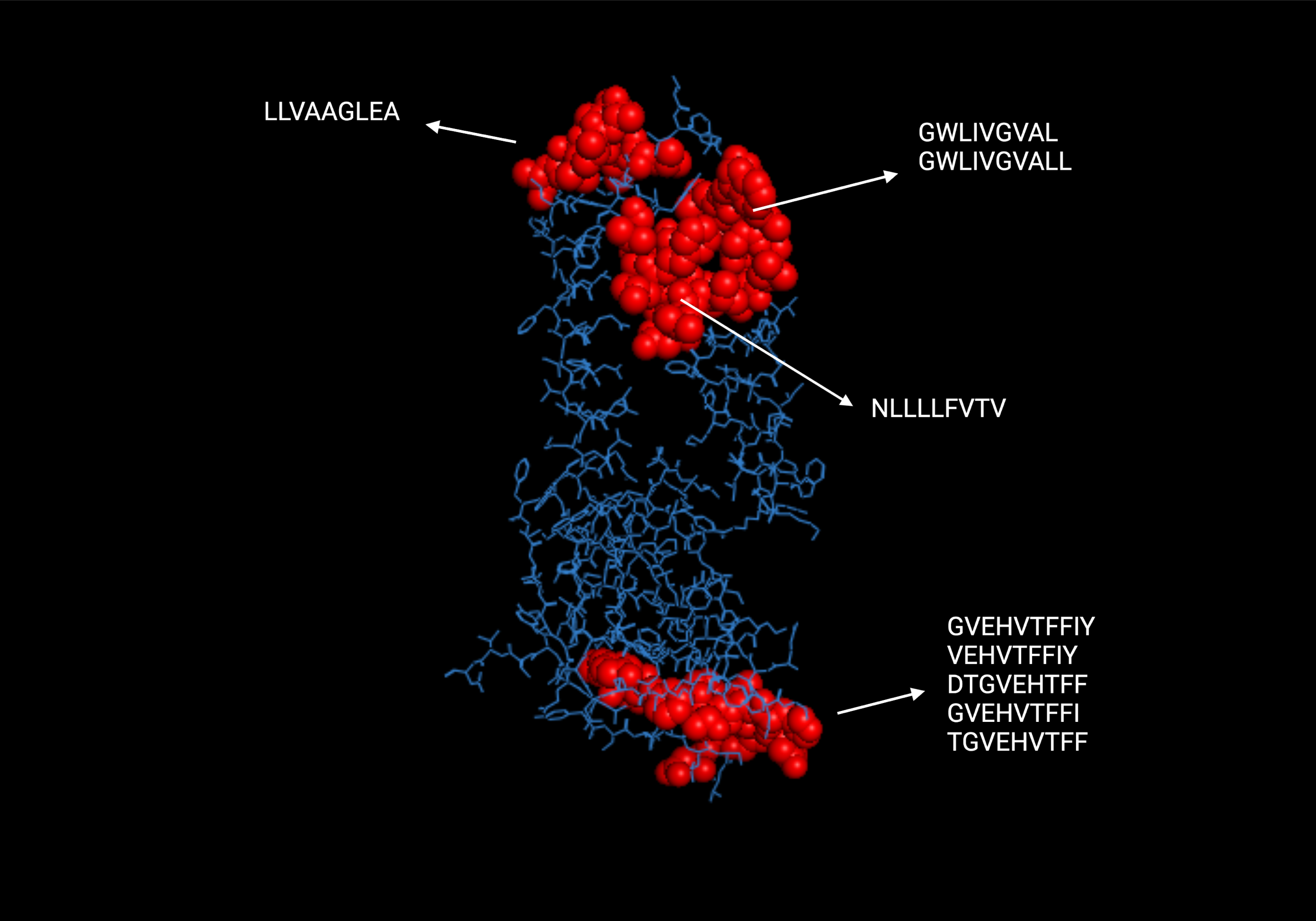

**Supplementary Figure 1a**. The positions of predicted CD8+ T cell epitopes (red) in 3D structure of reference ORF3a protein (blue).

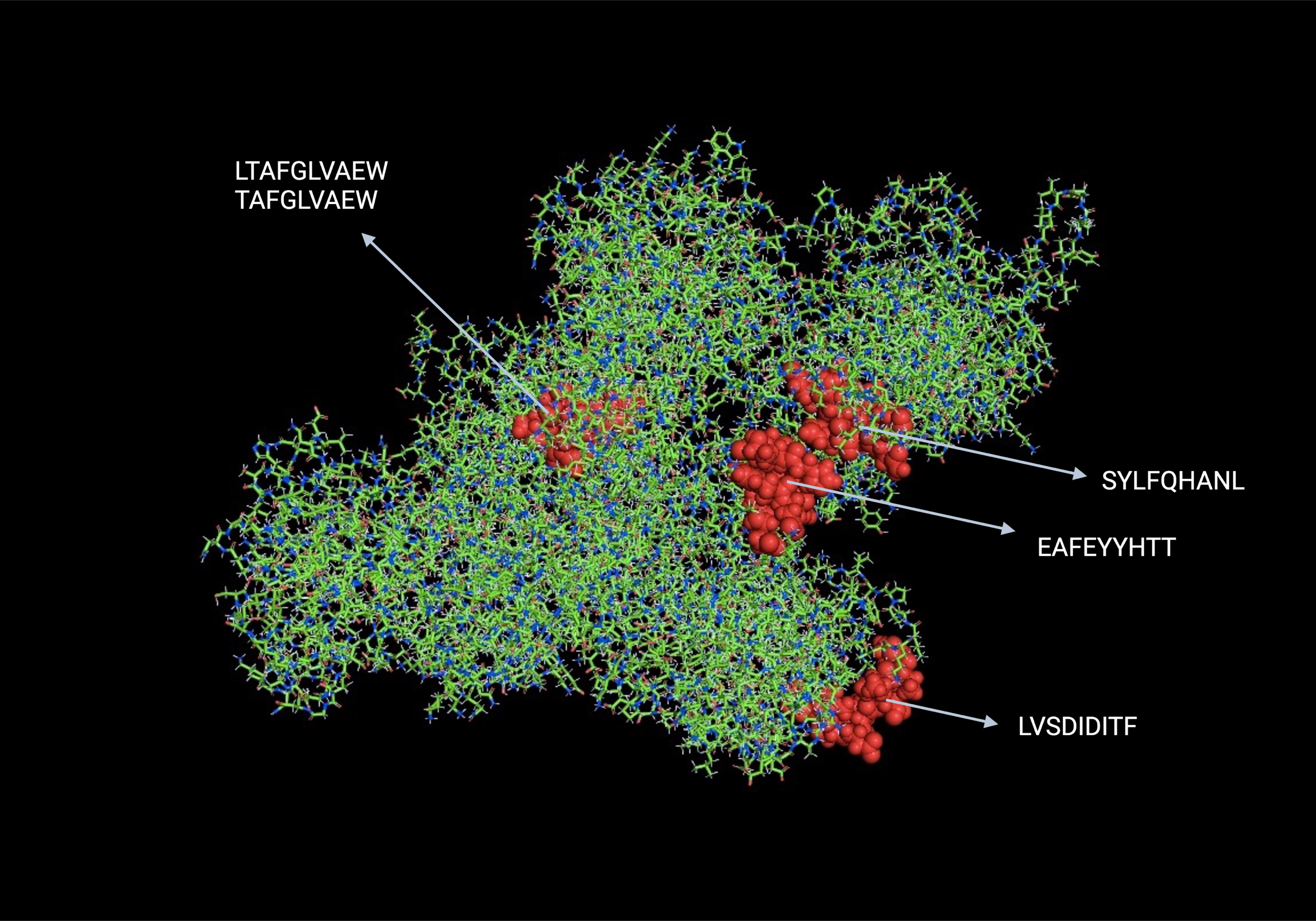

**Supplementary Figure 1b**. The positions of predicted CD8+ T cell epitopes (red) in 3D structure of reference nsp3 protein (green and blue)

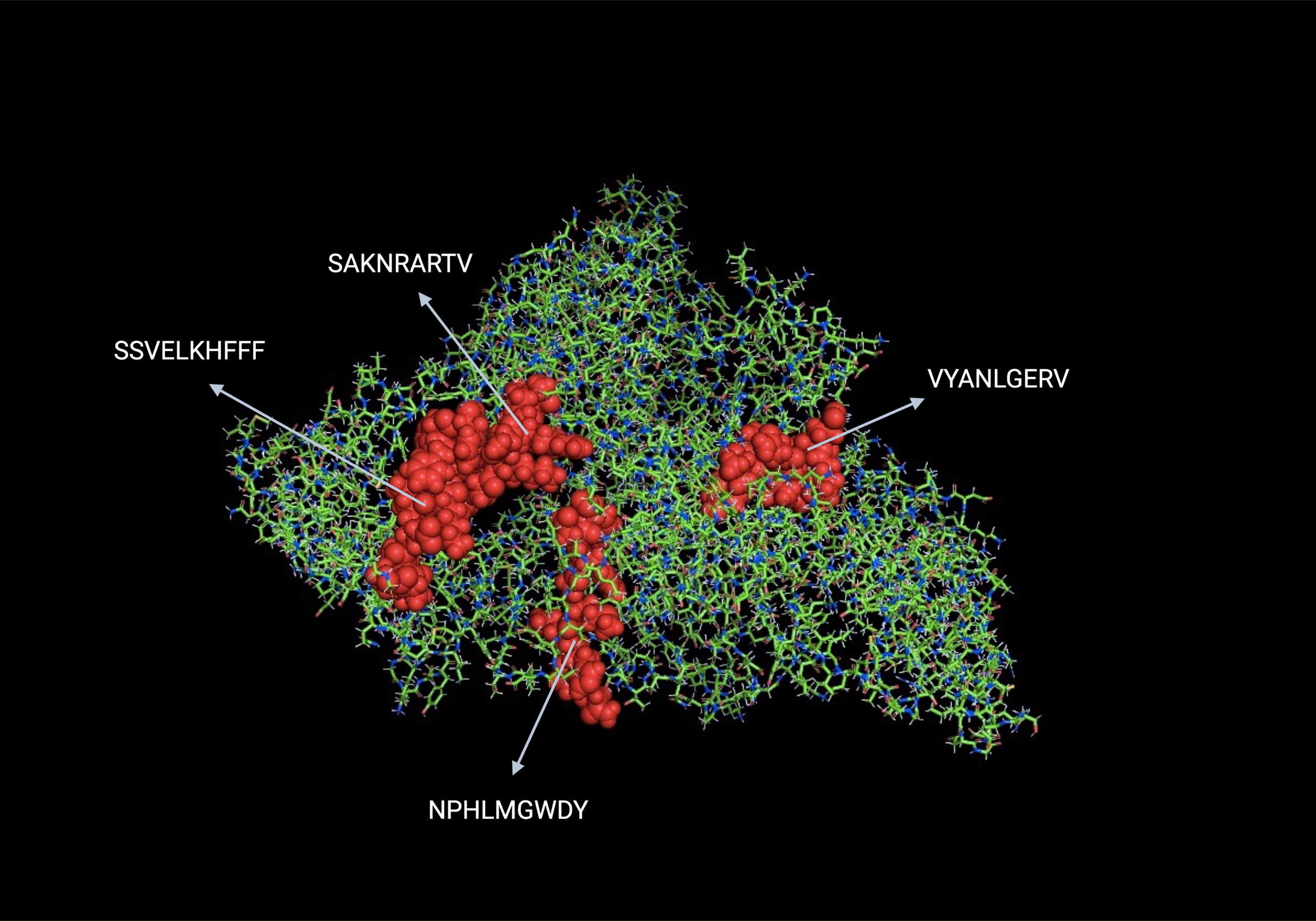

**Supplementary Figure 1c**. The positions of predicted CD8+ T cell epitopes (red) in 3D structure of reference nsp12 protein (green and blue)

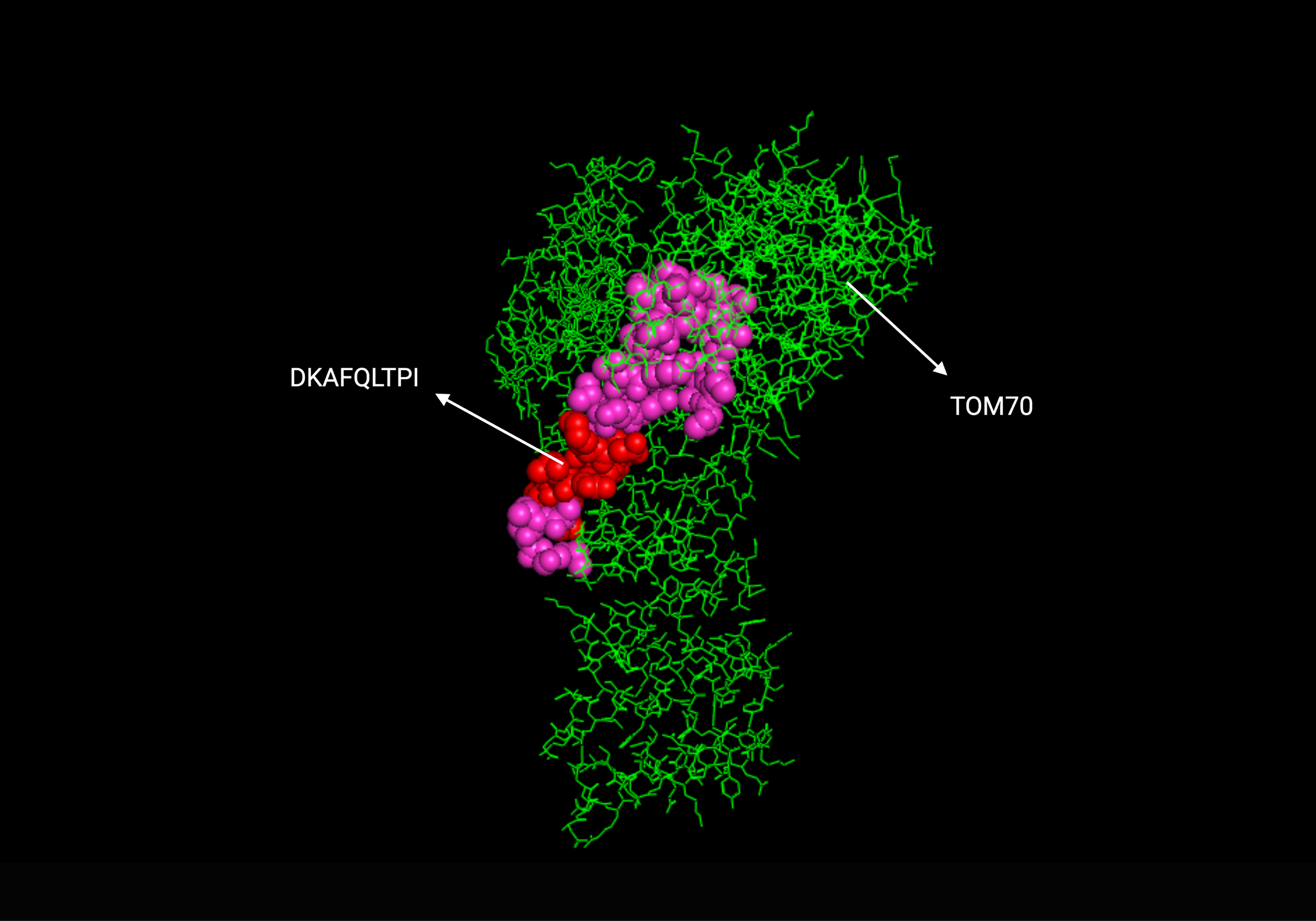

**Supplementary Figure 1d**. The position of the top epitope (red) of ORF9b (pink) in 3D structure and its interaction with TOM70 (green)
