## Supplementary Figure 2 for "Design of Peptide Vaccine for COVID19: CD8+ and CD4+ T cell epitopes from SARS-CoV-2 open-reading-frame protein variants"

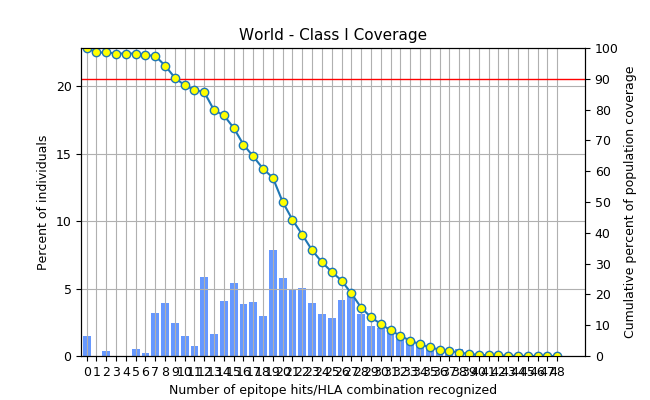

**Figure 2a**. World population coverage of top common immunogenic CD8 epitopes across VOC in ORF3a, excluding IFN-y filter. The population coverage was found to be 98.55%.

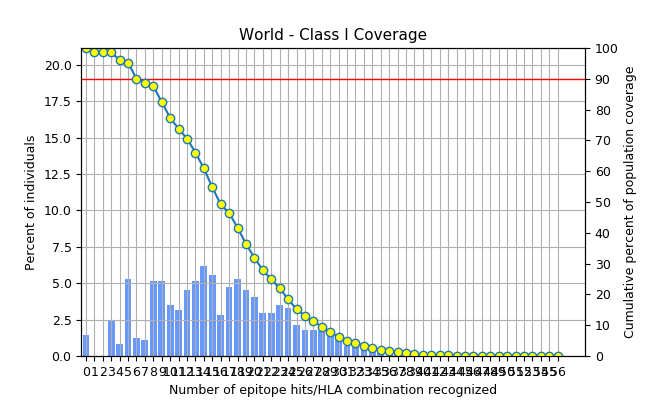

**Figure 2b**. World population coverage of top common immunogenic CD8 epitopes across VOC in nsp3, excluding IFN-y filter. The population coverage was found to be 98.55%.

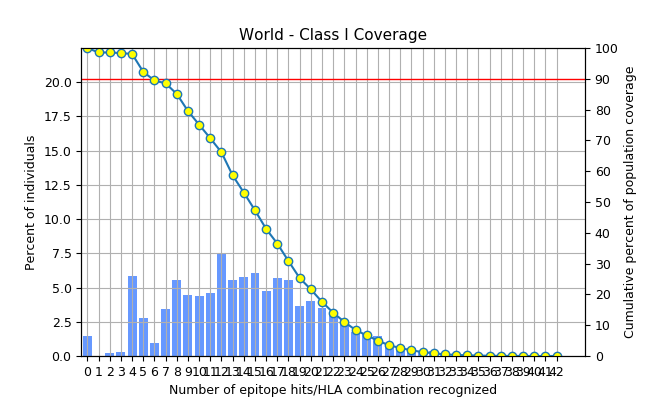

**Figure 2c**. World population coverage of top common immunogenic CD8 epitopes across VOC in nsp12, excluding IFN-y filter. The population coverage was found to be 98.55%.

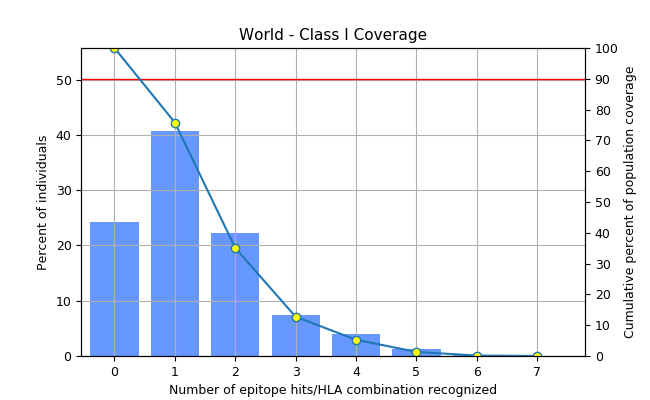

**Figure 2d**. World population coverage of top immunogenic CD8 epitopes in ORF9b, excluding IFN-y filter. The world population coverage was found to be 75.8%.

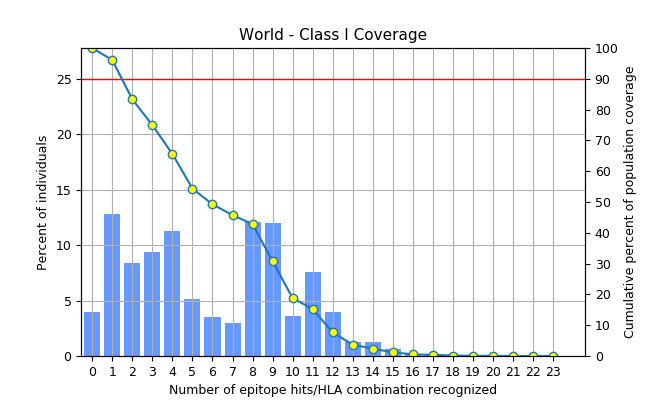

**Figure 2e**. World population coverage of top common immunogenic IFN-y inducing CD8 epitopes across VOC in ORF3a. The world population coverage was found to be 96.07%.

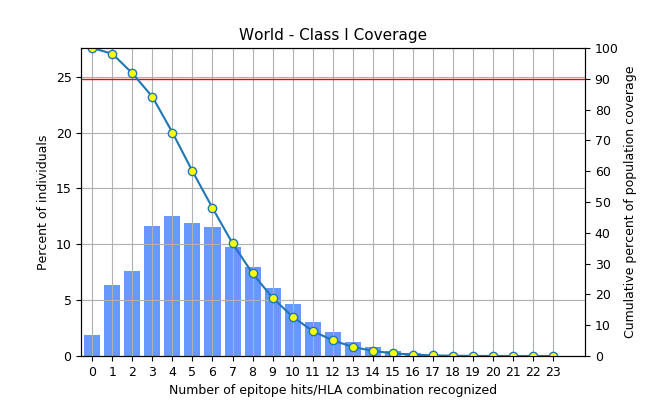

**Figure 2f**. World population coverage of top common immunogenic IFN-y inducing CD8 epitopes across VOC in nsp3. The world population coverage was found to be 98.12%.

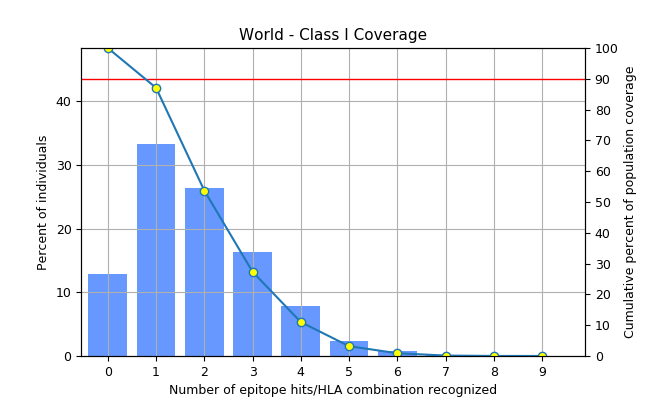

**Figure 2g**. World population coverage of top common immunogenic IFN-y inducing CD8 epitopes across VOC in nsp12. The world population coverage was found to be 87.08%.

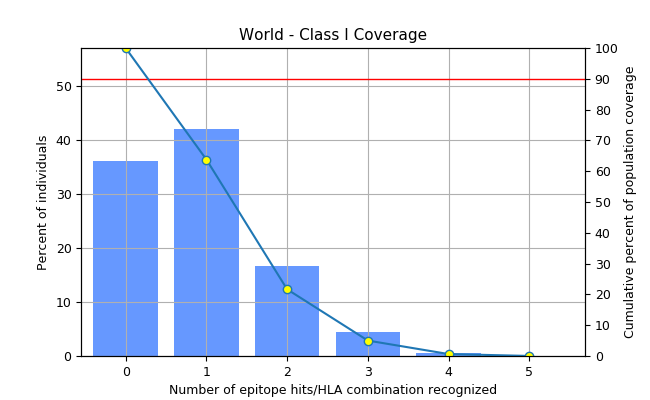

**Figure 2h**. World population coverage of top immunogenic IFN-y inducing CD8 epitopes in ORF9b. The world population coverage was found to be 63.8%.

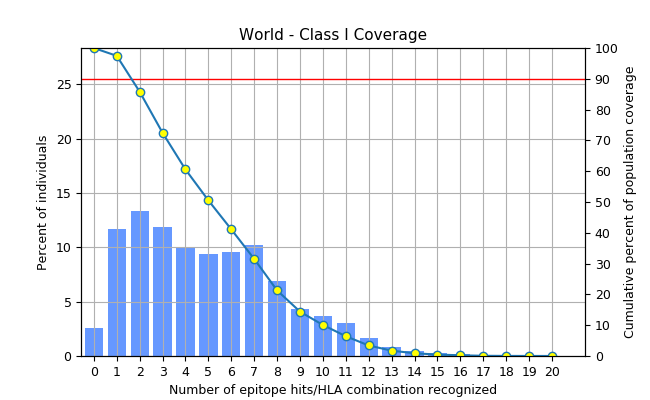

**Figure 2i**. World population coverage of top immunogenic CD8 epitopes across VOC in ORF3a with common mouse H2 allele restriction. 6 immunogenic CD8 epitopes were either weak or strong binders to murine H2 alleles. To increase population coverage, 2 top immunogenic CD8 epitopes from ORF3a were additionally obtained. The population coverage was found to be 97.45%.

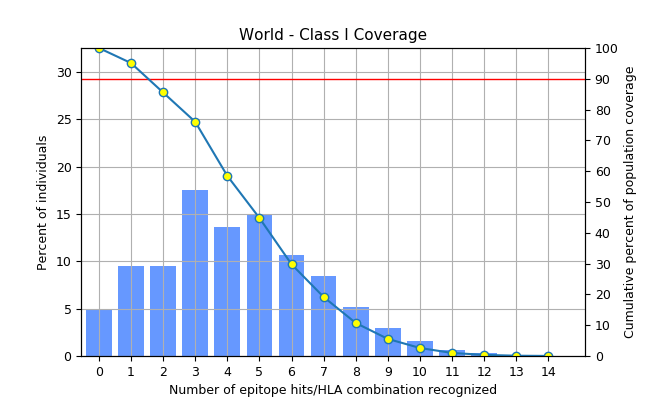

**Figure 2j**. World population coverage of top immunogenic CD8 epitopes across VOC in nsp3 with common mouse H2 allele restriction. The top 5 CD8 epitopes were predicted to protect 95.11% of the population.

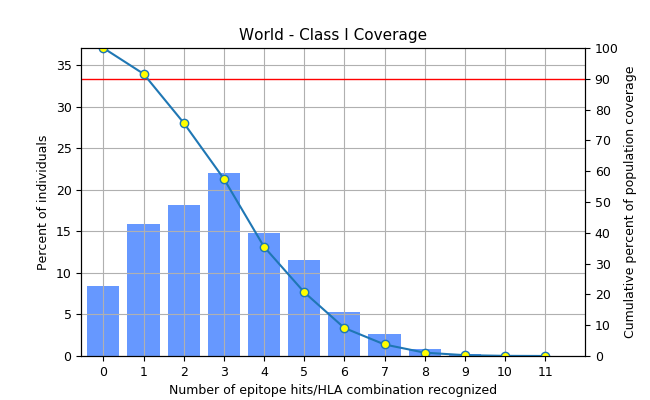

**Figure 2k**. World population coverage of top immunogenic CD8 epitopes across VOC in nsp12 with common mouse H2 allele restriction. The top 5 CD8 epitopes were predicted to protect 91.62% of the population.

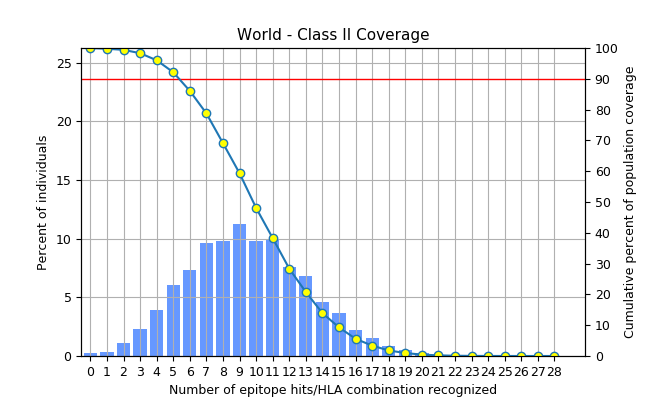

**Figure 2l**. World population coverage of top immunogenic IFN-y inducing CD4 epitopes across VOC in ORF3a. The population coverage of top 7 CD4 epitopes was found to protect 99.76% of the population.

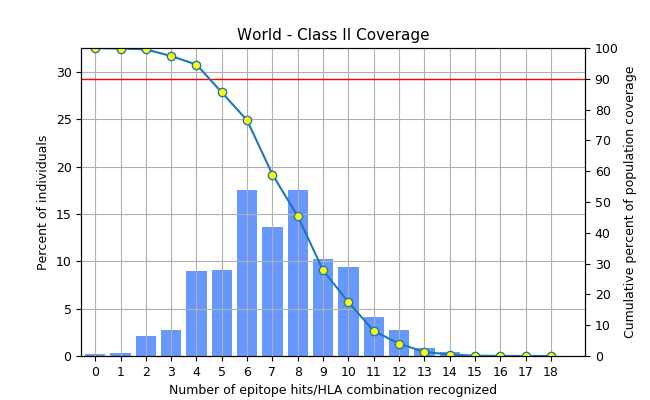

**Figure 2m**. World population coverage of top immunogenic IFN-y inducing CD4 epitopes across VOC in nsp3. 3 top immunogenic epitopes were predicted to elicit an immune response that covers 99.82% of the population.

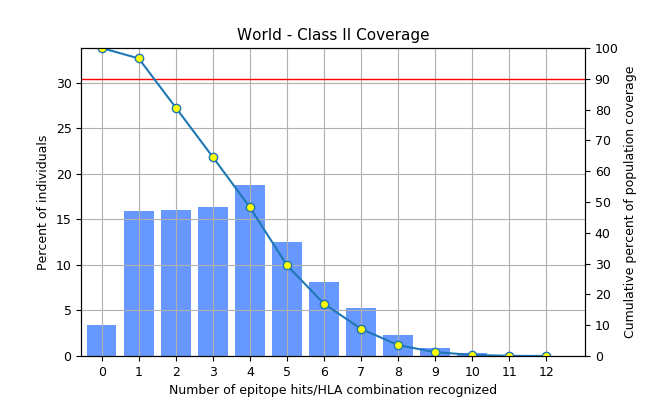

**Figure 2n**. World population coverage of top immunogenic IFN-y inducing CD4 epitopes across VOC in nsp12. The population coverage of top 6 CD4 epitopes was found to protect 96.62% of the population.

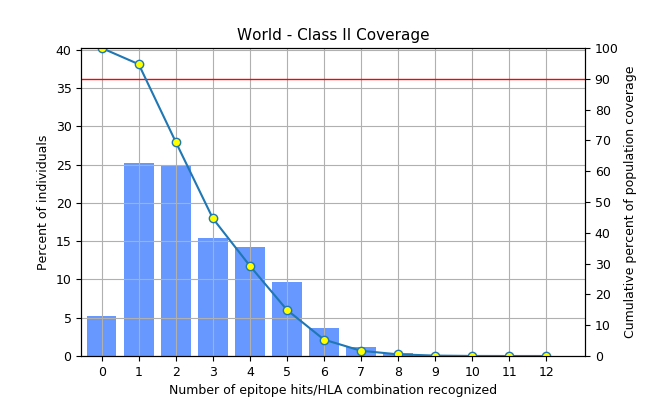

**Figure 2o**. World population coverage of top immunogenic IFN-y inducing CD4 epitopes in ORF9b. 4 top immunogenic CD4 epitopes were found to cover 94.74% of the population.

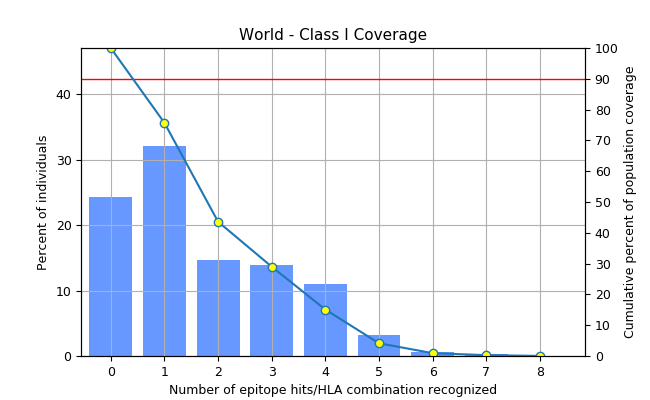

**Figure 2p**. World population coverage of top 3 CD8 epitopes from each protein: GWLIVGVAL (ORF3a), TAFGLVAEW (nsp3) and VYANLGERV (nsp12). These CD8 epitopes are highly immunogenic, IFN-y inducing, antigenic, non-allergenic, non-toxic, stable and common across all four variants of concern. The population coverage of these epitopes was predicted to be 75.72%.

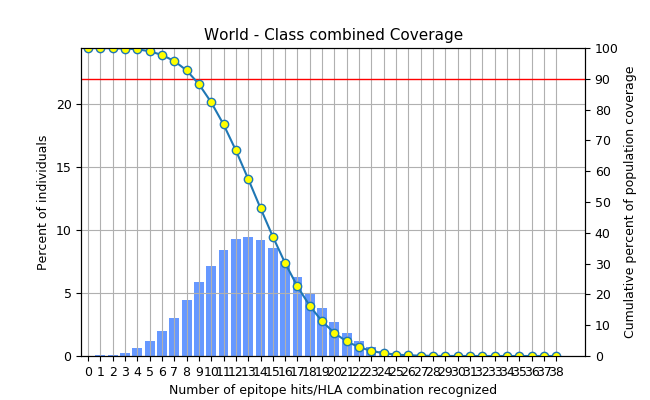

**Figure 2q**. World population coverage of top 3 CD8 epitopes: GWLIVGVAL (ORF3a), TAFGLVAEW (nsp3) and VYANLGERV (nsp12); and 6 CD4 epitopes: TAFGLVAEWFLAYIL (nsp3), LTAFGLVAEWFLAYI (nsp3), DLTAFGLVAEWFLAY (nsp3), PDILRVYANLGERVR (nsp12), RVYANLGERVRQALL (nsp12) and PFGWLIVGVALLAVF (ORF3a). These CD8 and CD4 epitopes are highly immunogenic, IFN-y inducing, antigenic, non-allergenic, non-toxic, stable and common across all four variants of concern. The population coverage of these epitopes is 99.99%.
